# A Perturbation Cell Atlas of Human Induced Pluripotent Stem Cells

**DOI:** 10.1101/2024.11.03.621734

**Authors:** Sami Nourreddine, Yesh Doctor, Amir Dailamy, Antoine Forget, Yi-Hung Lee, Becky Chinn, Hammza Khaliq, Benjamin Polacco, Monita Muralidharan, Emily Pan, Yifan Zhang, Alina Sigaeva, Jan Niklas Hansen, Jiahao Gao, Jillian A. Parker, Kirsten Obernier, Timothy Clark, Jake Y. Chen, Christian Metallo, Emma Lundberg, Trey Ideker, Nevan Krogan, Prashant Mali

**Affiliations:** Department of Bioengineering, University of California San Diego, CA, USA; Quantitative Biosciences Institute (QBI), University of California San Francisco, CA, USA; Gladstone Institute of Data Science and Biotechnology, J. David Gladstone Institutes, San Francisco, CA, USA; Department of Cellular and Molecular Pharmacology, University of California San Francisco, CA, USA; School of Medicine, University of California San Diego, CA, USA; Department of Medicine, University of Virginia, VA, USA; Department of Computer Science, The University of Alabama at Birmingham, VA, USA; Molecular and Cell Biology Laboratory, The Salk Institute for Biological Studies, CA, USA; Department of Bioengineering, Stanford University, CA, USA; Department of Pathology, Stanford University, CA, USA; Department of Bioengineering and Therapeutics Sciences, University of California San Francisco, CA, USA; Division of Cellular and Clinical Proteomics, Department of Protein Science, SciLifeLab, KTH Royal Institute of Technology, Stockholm, Sweden

**Author notes:** These authors contributed equally (Sami Nourreddine, Yesh Doctor, Amir Dailamy, Antoine Forget).

## Abstract

Towards comprehensively investigating the genotype-phenotype relationships governing the human pluripotent stem cell state, we generated an expressed genome-scale CRISPRi Perturbation Cell Atlas in KOLF2.1J human induced pluripotent stem cells (hiPSCs) mapping transcriptional and fitness phenotypes associated with 11,739 targeted genes. Using the transcriptional phenotypes, we created a minimum distortion embedding map of the pluripotent state, demonstrating rich recapitulation of protein complexes, such as strong co-clustering of MRPL, BAF, SAGA, and Ragulator family members. Additionally, we uncovered transcriptional regulators that are uncoupled from cell fitness, discovering potential novel pluripotency (JOSD1, RNF7) and metabolic factors (ZBTB41). We validated these findings via phenotypic, protein-interaction, and metabolic tracing assays. Finally, we propose a contrastive human-cell engineering framework (CHEF), a machine learning architecture that learns from perturbation cell atlases to predict perturbation recipes that achieve desired transcriptional states. Taken together, our study presents a comprehensive resource for interrogating the regulatory networks governing pluripotency.

## INTRODUCTION

Mapping genotype-phenotype relationships is a fundamental goal of cell biology, providing critical insights into how genetic elements influence cellular function. Characterization of this relationship for each cell type is essential for understanding development, disease mechanisms, and informing the engineering of cell state. A cell type of particular interest is human induced pluripotent stem cells (hiPSCs), which possess the remarkable ability to differentiate into any cell type, making them invaluable for disease modeling and regenerative medicine applications^1–3^. Substantial progress has been made in identifying key transcription factors and signaling pathways that regulate pluripotency and enable direct reprogramming. For instance, the roles of POU5F1 (also known as Oct4), SOX2, and NANOG in maintaining stem cell pluripotency are well-documented^4^, while FOXA1/2/3, MYOD1, and NEUROD1 have been identified as factors initiating reprogramming into ecto-, meso-, and endoderm lineages respectively^5^. However, our understanding of pluripotency is far from complete, and the dynamic processes and gene interactions governing cell state transitions in hiPSCs are still not fully understood. We conservatively estimate that the functional role of less than 2000 genes have been deeply characterized within the stem cell context^6^, and hypothesize that there exist yet unknown pathways critical to stem cell regulation.

Perturb-Seq, a high-throughput method combining CRISPR-based genetic perturbations with single-cell RNA sequencing, offers a powerful approach to elucidate these unknown pathways^7–10^. By systematically knocking down or activating genes (perturbations) and quantifying the resultant changes in gene expression^11–14^ at single-cell resolution, researchers can map the functional relationships between genes and cell states at an unprecedented scale. The single cell nature allows screening of individual perturbations to be done in a pooled manner, allowing thousands of perturbations to be assayed in parallel^15^. However, Perturb-Seq is not without its challenges; for one, there is no standard agreed upon analysis methodology due to (among other reasons) (i) zero inflated gene expression distributions^16^, (ii) incomplete perturbation penetrance^17^, and (iii) challenges with algorithmic scaling^18^. Further, although a genome-wide perturbation cell atlas^19^ would be an invaluable resource for understanding the fundamental regulation of a cell type of interest, there has to date only been one published study at the pan-genome scale (covering 9,867 expressed genes in K562 lymphoblast cells^15^), likely due to high financial cost and technical difficulty of genome-scale Perturb-Seq. Specifically, within hiPSCs the largest Perturb-Seq experiment to date covers only 27 genes^20^, though an analogous study in hESCs overexpressing a library of 1,836 transcription factors yielded strong insights into transcription factors biasing differentiation trajectories.^6^

We therefore foresee an expressed genome-scale perturbation cell atlas in hiPSCs as an invaluable resource for understanding pluripotency and informing rational cell state engineering. To this end, we generated a CRISPR interference (CRISPRi) Perturb-Seq dataset spanning the entire expressed genome in KOLF2.1J hiPSCs^21^, accompanied by a parallel fitness readout. To enhance the robustness of our findings, we validated key perturbations at multiple levels, including protein complex mapping through size-exclusion chromatography coupled to mass spectrometry (SEC-MS) and functional metabolic tracing using isotopologue analysis. These validation experiments confirmed that transcriptomic clusters derived from Perturb-Seq reflect interacting protein complexes and metabolic phenotypes in hiPSCs. Additionally, we developed a proof-of-concept machine learning architecture that predicts perturbations capable of driving cells towards desired transcriptional states, paving the way for rational engineering of cell states. Together, this dataset and its comprehensive validation provide a powerful resource for dissecting pluripotency regulation and exploring targeted cell state modifications.

## RESULTS

### Pilot Screens of a Panel of Chromatin Modifiers and Metabolic Enzymes CRISPRi Perturb-Seq in KOLF2.1J iPSCs

Our experimental and computational methodologies for the expressed-genome scale screen were first optimized using two pilot screens on a curated gene list (based on their known involvement in disease pathways) of 108 chromatin modifiers and 97 metabolic enzymes (**Supplementary Figure 1**). To perform genetic perturbations, we elected a CRISPRi strategy for the host of reasons outlined by Replogle et al.^15^, primarily that through scRNA-seq the extent of perturbation can be quantified for CRISPRi. We engineered a KOLF2.1J line that stably expresses KRAB^ZIM3^-dCas9^15,22^ (henceforth simply referred to as KOLF2.1Js; see Methods). For each gene, six sgRNAs were designed via a combination of selecting sgRNA designated by Replogle et al.^15^ and from CRISPick^23^ tool, which ranks gRNAs based on on-target efficiency and off-target effects, along with a total of 109 non-targeting control (NTC) sgRNAs. KOLF2.1J cells maintained in pluripotent maintenance media (mTeSR) were harvested at day 4 post-transduction and processed for single-cell RNA-sequencing (**Supplementary Figure 1A**). We confirmed robust expression of pluripotency factors NANOG and POU5F1 via immunostaining (**Supplementary Figure 1A**). For the chromatin modifiers screen, we sequenced to a depth of median >20,000 unique molecular identifiers (UMIs)/cell (**Supplementary Figure 1B**). We loaded ∼100K cells/channel into four channels of a 10X HT 3’ chip, which yielded a final coverage of >100 single cells with a single sgRNA assigned for >90% of perturbations (**Supplementary Figure 1C**). The KRAB^ZIM3^-dCas9 CRISPRi repression system induced fairly reliable repression, with ∼70% of perturbation targets experiencing >30% repression (**Supplementary Figure 1D**). We observed that generally, at least one of the top 3 CRISPick candidate sgRNA induced sufficient repression, and therefore in the expressed genome-scale screen elected to use only three sgRNA per gene.

To identify strong perturbations (i.e., those that likely alter cell state relative to NTC), we developed a custom computational pipeline that aimed to balance resolution with computational runtime owing to the massive scale of the datasets (**Supplementary Figure 2**). See Methods for full details. Briefly, after isolating single, live cells harboring a single sgRNA (**Supplementary Figures 2A,B,C**), we isolate a population of NTC cells that: (i) harbor NTC sgRNA that do not induce a fitness impact, (ii) are present in all three experimental batches, and (iii) do not have a deviant transcriptional profile (**Supplementary Figure 2D**). These are designated as core NTC cells. We then isolate perturbing sgRNAs that: (i) induce at least 30% mean on-target knockdown (**Supplementary Figure 2E**), and (ii) induce a transcriptional profile more deviant than 75% of core NTC sgRNA from the same batch (**Supplementary Figure 2F**). Additionally, we filter for cells experiencing a transcriptional profile more deviant than 75% of core NTC cells from the same batch (**Supplementary Figure 2F**). After filtering for perturbations with at least 25 remaining cells, differential expression analysis was performed via DESeq2^24^ (adj. p < 0.05) using core NTCs from the same batch as the reference for each perturbation. (**Supplementary Figure 2G**). Perturbations with at least 10-25 differentially expressed genes (DEGs) were identified as strong perturbations. Finally, experimental batches were integrated by normalizing transcriptional profiles to core NTC cells within each batch (**Supplementary Figure 2H**). In the energy distance test, DNMT1 and UHRF1 were both identified as significant, demonstrated the largest energy distance, and were the #1/#6 perturbations inducing the most DEGs. Further, these two perturbations demonstrated co-clustering in a cell level UMAP projection (**Supplementary Figure 1I**), and were located adjacent in the MDE of this dataset (**Supplementary Figure 1J**). This result corroborates previous results, as UHRF1 is known to recruit DNMT1 for DNA methylation maintenance^25^, which plays a critical role in regulating gene expression in stem cells^26^. We additionally observed co-clustering of deacetylases, such as the class IIa HDACs HDAC4/HDAC7^27^ (**Supplementary Figure 1J**). These observations, in conjunction with the observed lack of co-clustering between perturbed and NTC cells in general (**Supplementary Figure 1I**), gave us confidence that our experimental and computational pipelines were suitable for a larger scale screen.

We further tried to optimize the yield of single-cells with a single sgRNA assigned by titrating the number of cells loaded per 10X channel from between 50K to 200K per channel in the pilot screen of 97 metabolic enzymes (**Supplementary Figure 1K/L**). We found that loading 200K cells per channel yields a high multiplet rate and poor yield of usable cells. Loading 150K cells per channel did yield the highest number of usable cells, though at lower efficiency than loading 100K cells. Thus, for the expressed genome-scale study we aimed to load between 115K-125K cells per channel (which was, at the time, in line with 10X’s recommended maximum loading). Analysis of the perturbations to metabolic enzymes revealed smaller overall effect sizes compared to the chromatin modifiers as evidenced by the reduced overall number of strong perturbations and number of DEGs (**Supplementary Figure 1M**). We did observe that perturbations to TGFBR2 induced the second largest number of DEGs. Considering that TGFβ signaling is critical for pluripotency maintenance^28^, we would anticipate knockdown to TGFBR2 to induce a divergent cell state from wild type (WT) cells, which we observed at the cell level (**Supplementary Figure 1N**).

### Pan-Expressed Genome CRISPRi Perturb-Seq in KOLF2.1J iPSC

Building on our learnings from the pilot screen, we next scaled our efforts to generate a comprehensive Perturb-Seq dataset of KOLF2.1J hiPSC cells with an expressed genome-wide CRISPRi library targeting 11,739 unique genes (**Figure 1A**). For each gene, three sgRNAs were designed via CRISPick, along with a total of 478 non-targeting control sgRNAs. KOLF2.1J cells maintained in pluripotent maintenance media (mTeSR^29^) were harvested at day 6 post-transduction and processed for single-cell RNA-sequencing (**Figure 1A**). A total of ∼11 million cells were processed. Post filtering, our approach yielded more than 2.5 million single cells, each harboring one sgRNA, with a median of over 5,000 UMIs per cell (**Figure 1C**). On average, 88% of perturbations had a coverage of over 100 associated cells (**Figure 1D**), and 81% achieved greater than 30% mean target knockdown efficiency as assessed by expression of target transcript from scRNA-seq (**Figure 1E**).

**Figure 1:**
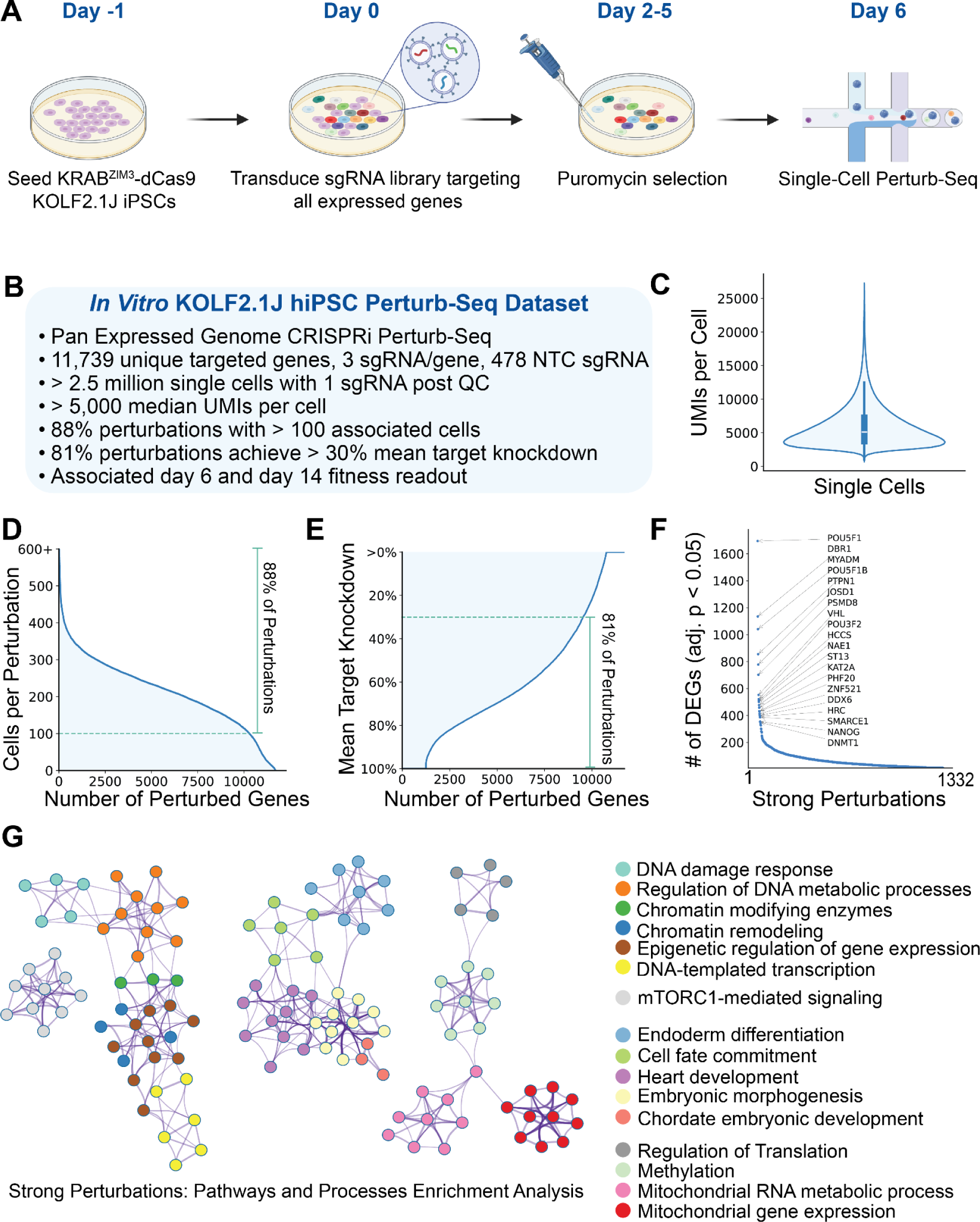
Pan-Expressed Genome CRISPRi Perturb-Seq in KOLF2.1J iPSC. **(A)** Experimental workflow of the in vitro CRISPRi Perturb-Seq in KRAB^ZIM3^-dCas9 KOLF2.1J hiPSCs. Cells were seeded on day −1, transduced with an sgRNA library targeting all expressed genes on day 0, selected with puromycin from day 2 to day 5, and harvested for single-cell Perturb-Seq on day 6. **(B)** Overview of the KOLF2.1J hiPSC Perturb-Seq dataset. This genome-scale CRISPRi dataset covers 11,739 unique targeted genes with 3 sgRNAs per gene, along with 478 non-targeting control (NTC) sgRNAs. The dataset includes >2.5 million single cells with a median of >5,000 UMIs per cell, 88% of perturbations have >100 associated cells, and 81% achieved >30% mean target knockdown efficiency. **(C)** Distribution of unique molecular identifiers (UMIs) per cell across single-cell samples, showing a robust capture of transcriptomic data with a median UMI count of >5,000 per cell. **(D)** Number of cells per gene across the library, highlighting that 88% of perturbations have >100 associated cells, ensuring sufficient statistical power for downstream analysis. **(E)** Distribution of mean target knockdown efficiency, indicating that 81% of perturbations achieve >30% knockdown of their target genes, confirming the efficacy of the CRISPRi system in KOLF2.1J cells. **(F)** Top perturbations by number of differentially expressed genes (DEGs) with adjusted p-value < 0.05. Known pluripotency regulators, such as POU5F1, NANOG, and PSMD8, rank among the strongest perturbations. **(G)** Functional network of strong perturbations computed with Metascape. Each cluster represents key biological processes, and each node represents a gene, color-coded by function.

Following computational analyses (**Supplementary Figure 2**), we identified 1,332 strong perturbations (**Figure 1F**); perturbations with at least 10 DEGs were classified as strong perturbations. The strongest perturbations identified in the screen highlight well-established pluripotency or developmental factors, such as POU5F1^4^, NANOG^30^, PTPN1 (also known as PTP1B)^31^, ZNF521 (also known as EZHF or evi3)^32,33^, PSMD8^34^, and SMARCE1^35^, confirming their critical roles in maintaining the pluripotent state of iPSCs. To further understand the functional impact of these strong perturbations, we performed pathway enrichment analysis on them using the Metascape tool^36^ (**Figure 1G**). The network plot reveals clusters of enriched pathways and processes, which include chromatin modifying enzymes (Log10(p-value)=-16.64), DNA damage response (Log10(p-value)=-11.28), and mTORC1-mediated signaling (Log10(p-value)=-8.79), among others. Notably, pathways related to embryonic morphogenesis (Log10(p-value)=-10.99), cell fate commitment (Log10(p-value)=-9.66), endoderm differentiation (Log10(p-value)=-9.81), and mitochondrial metabolic processes (Log10(p-value)=-12.17) were identified. All together, this observation highlights the robustness of the screen in generally capturing key regulators essential for pluripotency, validating the central role of these genes in the self-renewal and pluripotency network of hiPSCs.

### Functional gene inference through analysis of transcriptional signatures in KOLF2.1J

For the 1,332 strong perturbations, we computed the pairwise Pearson correlation matrix using the top 2,000 highly variable genes as features (**Figure 2A**). The presence of distinct clusters across the diagonal suggests that certain gene perturbations have highly correlated transcriptional outcomes, indicating potential involvement in related biological processes or pathways. The correlation matrix was then embedded into a Minimal Distortion Embedding (MDE) map^37^, resulting in the organization of these perturbations into 50 distinct clusters identified by Leiden clustering (**Figure 2C**, **Supplemental Figure 1I**). Functional enrichment analysis was then performed on these MDE-derived clusters via gProfiler^38^ and used to label the function of the perturbation clusters based on CORUM complexes and Gene Ontology terms. We observed an enrichment of protein complexes and biological pathways within the MDE-derived clusters compared to random pairs, suggesting that biologically meaningful relationships between perturbations were effectively captured (**Figure 2B**). Cellular transcriptional profiles were additionally visualized using UMAP, illustrating how perturbations induce diverse cellular states (**Figure 2D**). Leiden clustering identified distinct transcriptional clusters, suggesting that perturbations drive cells into unique transcriptional identities. When comparing NTC cells, which are densely concentrated in a central cluster, to perturbed cells, which are more widely dispersed, it became evident that the identified strong perturbations introduced substantial transcriptional changes. We then visualized the complete network of regulatory interactions between all strong perturbations (**Supplementary Figure 3A**). Each strong perturbation was represented by a node, with outgoing edges to other strong perturbations that it regulates (i.e., an edge from A to B implies B is a DEG of A). Using PageRank^39^ on the reversed graph, we prioritized the influence of perturbations on the network. Put simply, nodes are given high importance for regulating many other nodes. Nodes that regulate nodes with high importance are also given higher importance. Upon visualization, we see a comprehensive picture of the regulatory interactions governing KOLF2.1J hiPSCs (**Supplementary Figure 3A**). When assessing the top 50 regulators (as determined by PageRank, **Supplementary Figure 3B**), we identify several core pluripotency factors, such as POU5F1 and NANOG, thatare essential for maintaining stem cell identity, differentiation effectors, such as PTPN1 and ZNF521, and key chromatin modifiers, such as SMARCE1 and ARID2, supporting their relevance in stem cell regulation^31,32,35,40–44^. We then isolated nodes associated with perturbations in pluripotency related clusters in the MDE (Clusters 31, 38, and 46, **Figure 2C and Supplementary Figure 4A)** and visualized the regulatory relationships between them (**Supplementary Figure 3C).** We observed that a large number of perturbations in these clusters regulate both POU5F1 and ZNF521 expression, placing them at the center of the pluripotency regulatory network, reinforcing their critical role in stem cell maintenance^32,43–46^.

**Figure 2:**
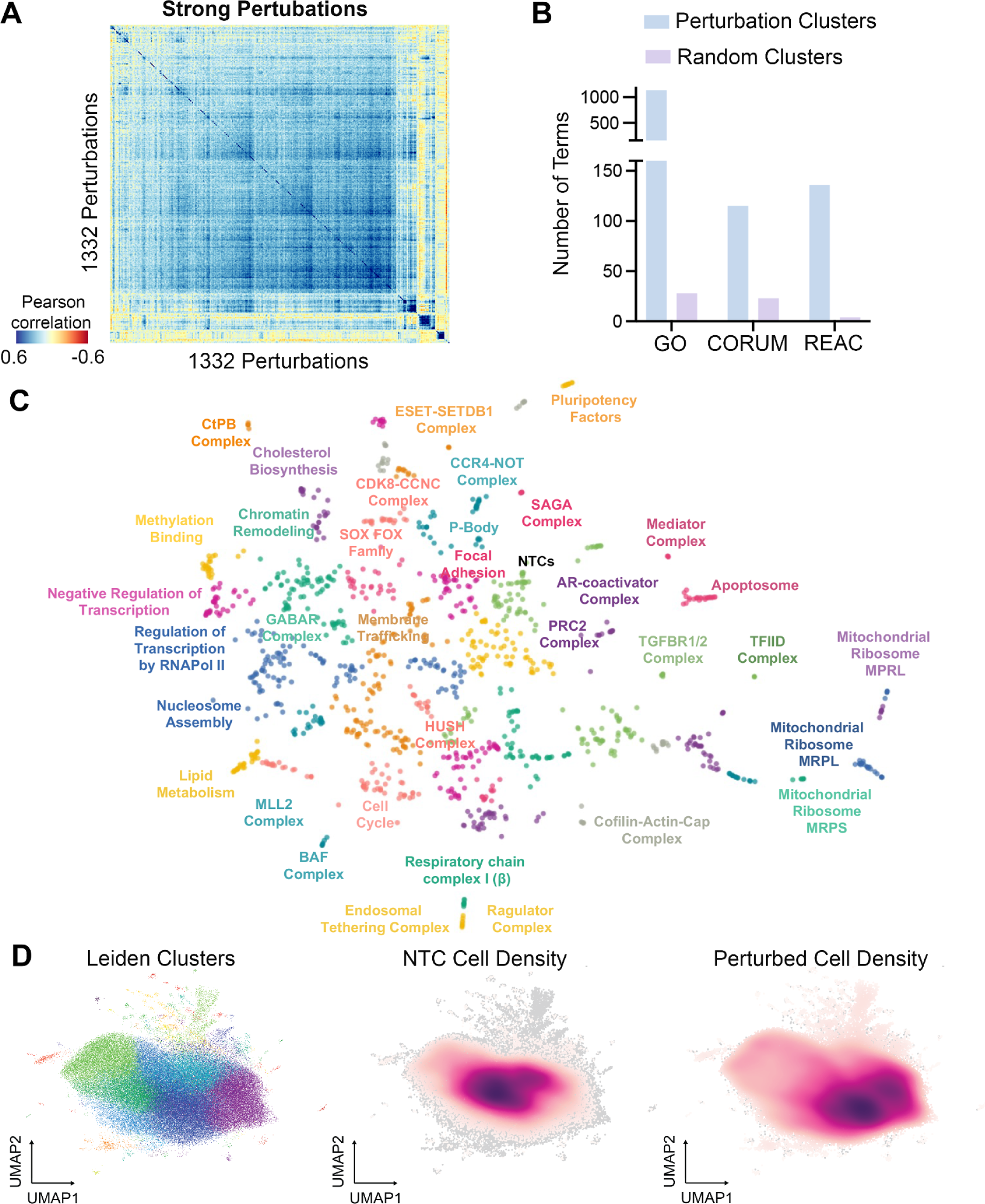
Correlation Landscape of Strong Perturbations. **(A)** Pearson correlation clustermap of pseudo bulked transcriptional profiles of strong perturbations **(B)** Functional enrichment analysis comparing the number of terms identified in the 50 perturbation clusters derived from the Minimal distortion embedding (C) versus 50 clusters with randomized perturbations across three databases: Gene Ontology (GO), CORUM, and Reactome (REAC). **(C)** Minimal distortion embedding (MDE) of pseudo bulked transcriptional profiles of strong perturbations. Clusters are labeled with associated enriched GO/CORUM/GSAI terms. **(D)** UMAP plots of cell density distributions: (left) Leiden clusters showing the organization of perturbations, (center) cell density distribution of non-targeting controls (NTCs), and (right) density distribution of perturbed cells.

### Comprehensive Fitness Profiling in KOLF2.1J

While transcriptional profiling provides valuable insights into the functional relationships and regulatory networks of genes **(Figure 2**), it requires capturing a sufficient number of cells to reliably profile the transcriptome. As such, this approach is inherently biased, and can potentially miss perturbations that have a profound negative fitness impact, as corresponding perturbations would result in significant cell loss. To address this limitation and gain a more comprehensive understanding of gene essentiality and functional dependencies in iPSCs, we in tandem with Perturb-Seq performed a pan expressed genome fitness screen in KOLF2.1Js cell line. Genome scale CRISPR fitness screens have previously been shown to give rich insights into genes serving as critical regulators.^47–52^ Using the same cell population that was assayed in the CRISPRi expressed genome-scale Perturb-Seq study, bulk genomic DNA was harvested on day 6 and further on day 14 to assess changes in sgRNA abundance, which reflects the impact of gene knockdown on cell fitness (**Figure 3A**). The fitness impact of each gene was determined by combining the gRNA enrichment and depletion data to obtain a gene-level fitness effect. Over time, there was a noticeable change in the number of genes exhibiting significant changes in fitness. At day 6, 1,067 genes were identified as significantly depleted (fitness z-score < −1), which increased to 1,428 by day 14, indicating the progressive identification of essential genes. Similarly, the number of enriched genes (fitness z-score > 1) increased from 77 at day 6 to 204 at day 14 (**Figure 3B**). This gradual accumulation of both enriched and depleted genes over time suggests that certain gene perturbations may induce delayed effects on cellular fitness. It also indicates that prolonged screening durations can capture a broader spectrum of gene functions, as genes involved in slower cellular processes or long-term adaptations may only become apparent at later time points. We then performed a functional enrichment analysis of perturbations that are enriched and depleted at day 14 (**Figure 3C**), which revealed distinct biological processes associated with both the depleted and enriched gene sets. The genes identified as essential in KOLF2.1Js (shown in red) correspond to fundamental biological processes, including translation, cell cycle, and DNA replication, highlighting their critical roles in maintaining cellular viability and proliferation. In contrast, enriched perturbations exhibit a different functional profile, indicating that these genes are involved in chromatin remodeling (SAGA complex components) as well as pathways related to apoptosis and DNA damage response. This suggests that many of these genes may function as tumor suppressors, where their disruption could provide a fitness advantage to the cells. For instance, TP53, BAX, BAK1, PMAIP1, and BBC3 (also known as PUMA) are well-established apoptosis and DNA damage response genes that, when depleted, are known to enhance proliferation and survival in pluripotent stem cells^53–56^. The SAGA complex, in particular, has also been previously identified as an important regulator in pluripotent stem cells^57^. This highlights the critical role of chromatin remodeling and apoptosis pathways in modulating cellular fitness within iPSCs.

**Figure 3.**
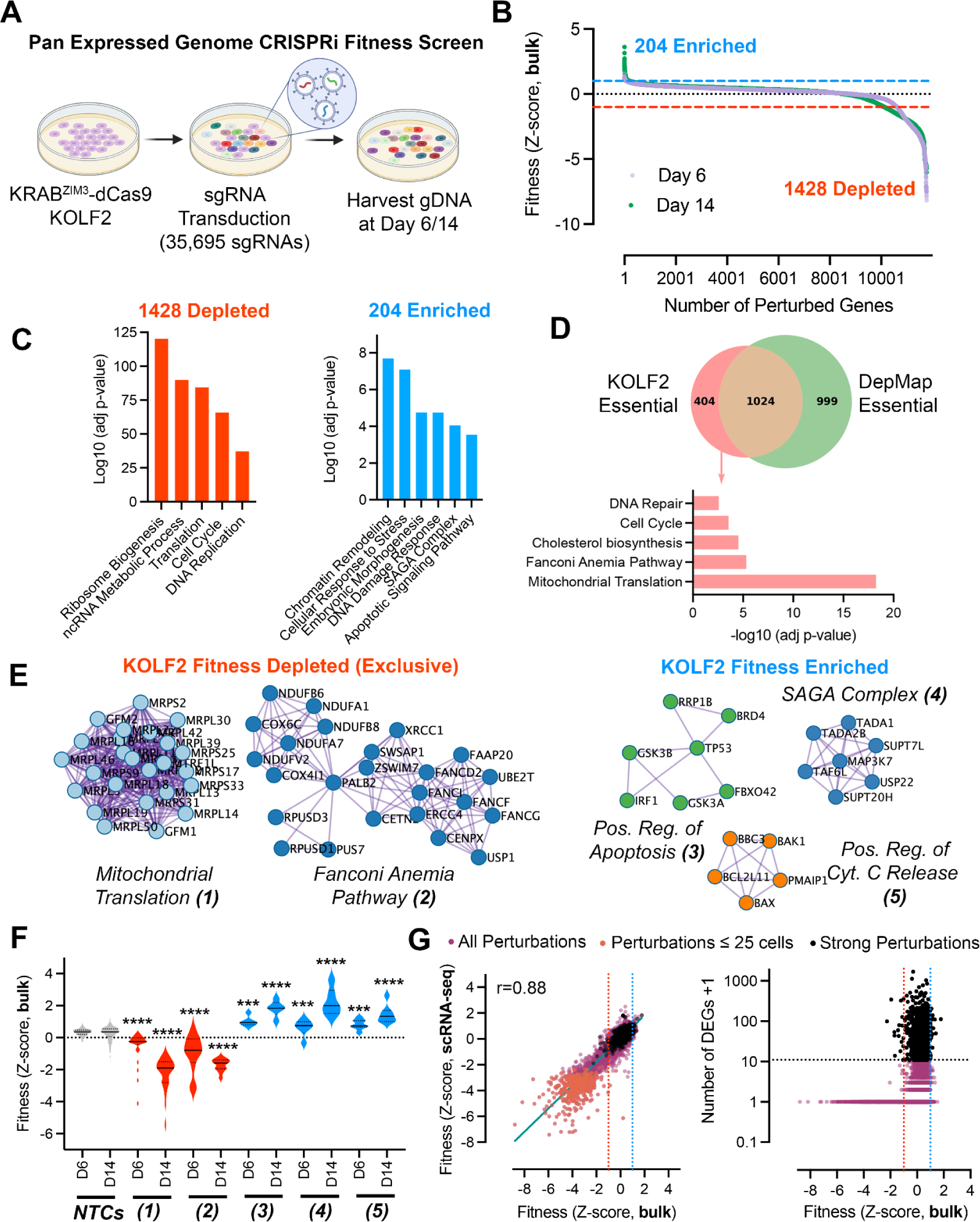
Undifferentiated hiPSC Fitness Screens. **(A)** Experimental workflow, KOLF2.1J constitutively expressing KRAB^ZIM3^-dCas9 were transduced with lentivirus libraries encoding for single guide RNA (sgRNA). Cells were cultured in pluripotency maintenance media mTesR and were harvested at day 6 and day 14 post-transduction. **(B)** Waterfall plots of Genes ranked by Z-score in descending order at days 6 and 14. Blue numbers correspond to enriched perturbations (Z-score>1;p-value<0.05), while red numbers correspond to depleted perturbations (Z-score<-1 p-value<0.05). **(C)** Functional enrichment analysis of the significant perturbations decreasing (red) or increasing fitness (blue) in KOLF2.1J. **(D)** Venn diagram representing overlap between the 1428 most depleted perturbations (Z-score<-1 p-value<0.05) in KOLF2.1J (KOLF2.1J essential) and the common essential genes identified in 1150 cell lines from DepMap portal (24Q2). In pink, functional enrichment analysis of the depleted perturbations that are exclusively found in KOLF2.1J. **(E)** Protein-Protein interaction enrichment analysis of the top depleted (red) and enriched (blue) perturbations. **(F)** Distribution of fitness Z-score at day 6 (D6) and day 14 (D14) of 478 unique NTC sgRNAs (NTCs) or Mitochondrial Translation 30 genes/90sgRNAs (1) Fanconi Anemia Pathway 12 genes/36sgRNAs(2) Positive Regulation of Apoptosis 7 genes/21sgRNAs (3) SAGA Complex 6 genes/18sgRNAs(4) Positive Regulation of Cytochrome C Release 5 genes/15sgRNAs(5). Asterisks indicate significant differences to D14 NTC condition (*p < 0.05, **p < 0.01, ***p < 0.001, ****p < 0.0001) **(G)** Left: Correlation between fitness calculated from bulk genomic DNA extraction at day 6 versus fitness calculated from abundance of cells captured by single-cell RNA-sequencing. Right: relationship between fitness (Z-score, bulk) and the number of differentially expressed genes (DEGs + 1) on a logarithmic scale. Each point represents a perturbation, points in purple represent all perturbations detected (11,688), in orange perturbations detected with less than 25 cells (427), in black the strong perturbations with more than 10 DEGs (1332).

We then compared our list of depleted (essential) genes in KOLF2.1Js to the essential gene list from the DepMap database (**Figure 3D**), which was generated from CRISPR screens across 1,150 cell lines, primarily consisting of cancer cells (Release 24Q2^47^). This comparison revealed that many of the essential genes identified in KOLF2.1Js overlap with the core set of essential genes found across diverse cell types, underscoring the identification of a fundamental group of genes crucial for cellular survival. However, a significant subset of genes that were exclusively essential in KOLF2.1Js iPSCs were also identified. Functional enrichment analysis of these KOLF2-exclusive essential genes showed notable enrichment in pathways such as mitochondrial translation (**Figure 3E**), indicating a heightened metabolic sensitivity in stem cells, and the Fanconi anemia pathway (**Figure 3E**), which underscores the high sensitivity of iPSCs to DNA damage and repair mechanisms^58–60^. We validated the effectiveness of our CRISPRi system by qPCR and observed strong and consistent repression of more than 24 genes including MRPL11 and MRPL14 (**Supplemental Figures S3C, S3D, S3E, S3F**). In addition, enrichment in the cholesterol biosynthesis pathway was also observed, corroborating the link between cholesterol metabolism and the maintenance or proliferation of pluripotent stem cells^61,62^. Using Metascape^36^ protein-protein interaction (PPI) enrichment tools, we then explored the PPI network of the most depleted and enriched perturbations, and found functional interdependencies among these genes, demonstrating that they not only share similar fitness impacts when perturbed but are also closely connected as known physical interactors within cellular pathways (**Figure 3E**). We then examined the fitness impact at day 6 and 14 corresponding to each of the PPI clusters identified compared to NTC (**Figure 3F**). NTC sgRNA abundance remained steady throughout the experiment, indicating minimal impact on cell fitness. In contrast, the clusters corresponding to mitochondrial translation and the Fanconi anemia pathway showed a progressive depletion at both day 6 and day 14, highlighting their essentiality for iPSC survival. Meanwhile, clusters related to the positive regulation of apoptosis and the SAGA complex exhibited progressive enrichment over time, suggesting that perturbations in these pathways confer a fitness advantage. In addition to the fitness impact observed at the functional cluster level, we also examined the fitness effects at the individual gene level within these clusters (**Supplemental Figures S3A, S3B**) providing detailed fitness z-scores for specific perturbations from the mitochondrial translation, Fanconi anemia, apoptosis regulation, and SAGA complex clusters.

We next compared the quantification of fitness between day 6 bulk genomic DNA extraction to cell abundance from guide RNA calling from the post-QC single-cell RNA-seq screen (**Figure 3G, left panel**). A strong correlation (r=0.88) between the fitness estimates obtained from both methods was observed, demonstrating the robustness of scRNA-seq in accurately assessing fitness and, similarly, validating the reliability of the bulk screen. However, it was also evident that perturbations with deep negative fitness impacts were often missed in the single-cell RNA-seq data, as these perturbations resulted in capturing fewer than 25 unique cells by day 6. We then investigated the correlation between the number of DEGs and fitness impact, and did not observe any significant correlation, which we hypothesize is due to the unique biology of pluripotent stem cells (PSCs) (**Figure 3G, right panel**). In PSCs, essential genes are often involved in maintaining core cellular functions including pluripotency, metabolism, and cell cycle regulation, where even slight perturbations can significantly impact cell fitness. However, recent insights suggest that the maintenance of pluripotent identity may be regulated independently from cell fitness^63^. While fitness screens traditionally emphasize survival and proliferation, they may overlook key pluripotency regulators that do not directly influence cell viability. This observation was also well illustrated in the recent Rosen et al. study^63^, highlighting that pluripotent identity and fitness may be governed by distinct mechanisms, with some genes contributing to the maintenance of the pluripotent state without impacting cell fitness. Together, these findings indicate that in pluripotent stem cells, fitness impact is more closely tied to disruptions in fundamental cellular machinery rather than the scale of transcriptional reprogramming. This reflects the sensitive and highly regulated nature of PSCs in balancing pluripotency and survival, where certain transcriptional regulators maintain pluripotency independently of fitness, expanding our understanding of the regulatory landscape of stem cell identity.

### KOLF2.1J Specific Perturbation Signatures and MDE Cluster Specific Fitness Impacts

To gain insights into cell type specific regulatory networks, the strong perturbations identified in our genome-scale KOLF2.1J Perturb-Seq study were compared with those observed in a K562 genome-scale Perturb-Seq experiment^15^. Our results indicate that strong perturbations are predominantly cell-type specific, with 1,069 unique to KOLF2.1J and 1,703 exclusive to K562, while only 263 perturbations are shared between the two cell types (**Figure 4A**). While we acknowledge the important caveat that the computational pipelines used to determine strong perturbations in the two studies are different, we anticipate that perturbations inducing strong transcriptional impacts will be generally similarly identified regardless of the pipeline used making these comparisons informative. When examining the KOLF2-exclusive strong perturbations, we observed that they are primarily involved in processes related to development and pluripotency (**Figure 4B**). Specifically, these perturbations are enriched in pathways such as cell fate commitment, chromatin remodeling, pluripotency regulation, transcriptional regulation of pluripotent stem cells, and germ layer formation, highlighting perturbations that are essential for maintaining and regulating the pluripotent state. This enrichment highlights the distinct developmental and differentiation potential of a pluripotent stem cell compared to a more lineage-restricted cell line, such as K562^64^.

**Figure 4.**
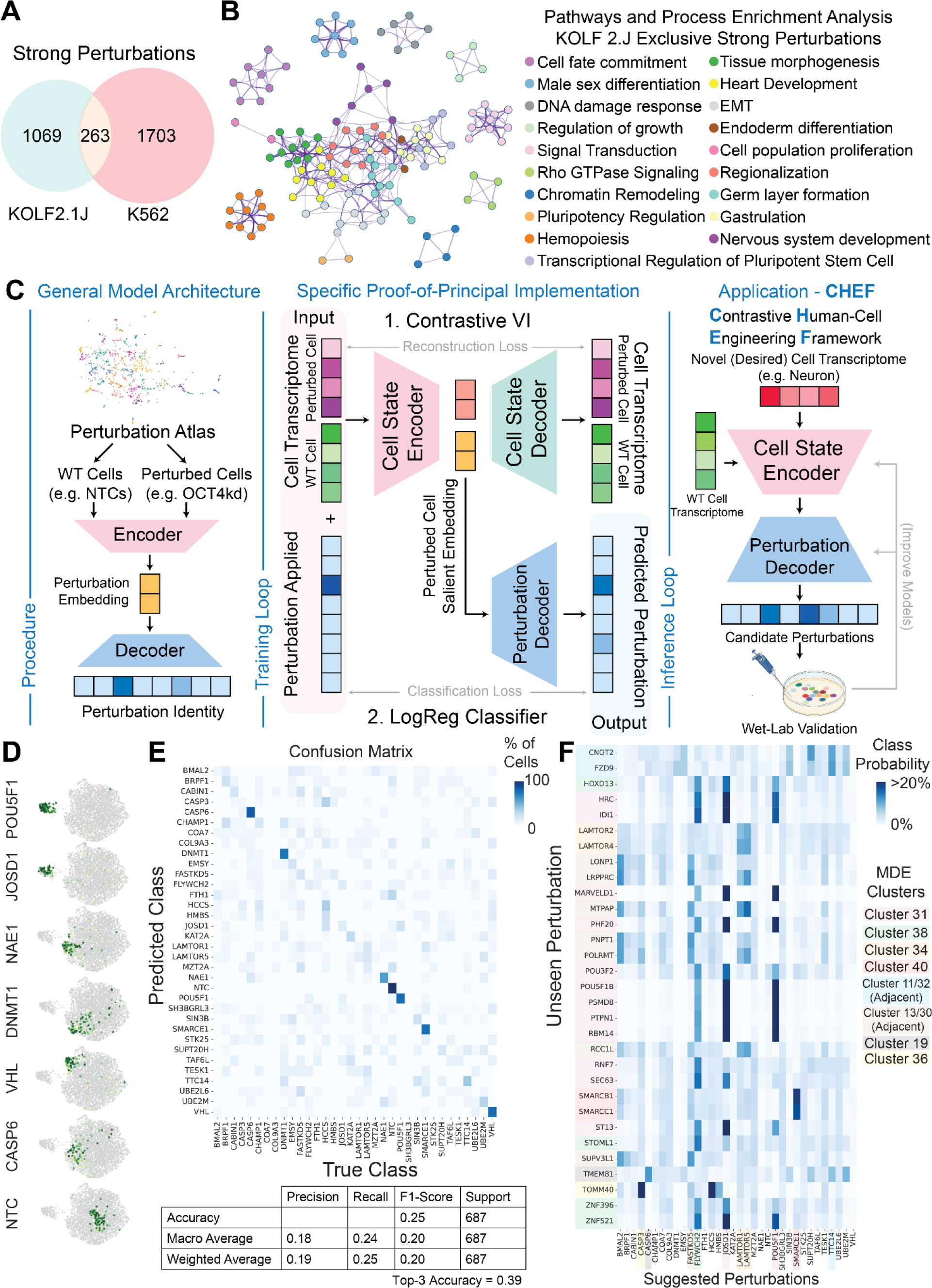
A Machine Learning Framework Towards Cell State Engineering. **(A)** Overlap between strong perturbations identified in KOLF2.1J (this study) and K562 (Replogle et al.^15^) datasets. **(B)** Metascape pathways and process enrichment analysis on strong perturbations identified exclusively in KOLF2.1J cells. **(C)** Proposed model architecture. Using perturbation atlas(es), a self- or semi-supervised encoder learns a perturbation embedding which corresponds to the difference in cell state between wild-type and perturbed cells. This embedding is then decoded by a supervised decoder to indicate which perturbation was applied. In our specific implementation, ContrastiveVI^73^ is first trained on the Perturb-Seq atlas to identify a salient embedding which isolates perturbation-specific effects. Then, a logistic regression (LR) classifier is trained on salient embeddings to decode them into the corresponding perturbation. Once this model is trained, a potential application is cell state engineering, where the transcriptome of a desired cell state (e.g. neuron) can be passed into the model, and it can infer candidate perturbations to make to WT cells to achieve that cell state. **(D)** UMAP visualization of salient embedding space for 6 perturbations + NTC **(E)** Confusion matrix and model metrics of LR classifier on predicting applied perturbations from salient cell state embeddings. **(F)** Inference of model on pseudo-bulked transcriptional profiles of previously unseen perturbations recapitulate observed clustering in MDE.

Using the perturbation-level MDE map (**Figure 2C**), a cluster-level representation was constructed where each node corresponds to the a cluster in the MDE, and the size of the node represents the number of member perturbations, retaining the overall layout observed in the original map (**Supplementary Figure 5A**). Edges are drawn between a node and its two nearest neighbors (as we observe neighbors tend to exhibit similar global transcriptional profiles, **Supplementary Figure 5B**), and each node is labeled with terms enriched using GO (Gene Ontology^65^), CORUM (Protein Complex Database^66^), and LLM GSAI (Large Language Model Gene Set Annotation and Interpretation^67^) tools, reflecting the main biological processes or cellular components associated with each cluster. Notably, Clusters 48, 42, 20, 46, and 34 are all related to mitochondrial function, and their connection on the MDE map underscores their close functional relationships, which aligns with their biological relevance in cellular energy production and regulation. Additionally, clusters representing mTORC signaling (Cluster 34), late endosome (cluster 34), and mitochondrial complex I assembly (MT Complex; Cluster 46) perturbations are located near each other, which is particularly interesting given the known role of mTORC being activated at late endosomes and promoting mitochondrial biogenesis^68^. Additionally, we found clusters related to Developmental Regulation (Cluster 38) and Pluripotency factors (Cluster 31), which are positioned nearby on the MDE map. Out of the 29 perturbations found in these two clusters, 15 rank among the top 30 strongest perturbations observed in the pan-expressed Perturb-Seq analysis (**Figure 1F**). In these two perturbations clusters, key transcription factors involved in pluripotency maintenance and neural development such as POU5F1^4^ and POU3F2^69^ were identified, as well as important developmental factors such as RBPJ^70^, PTPN1^31^, and ZNF521^32^. The perturbed cells from MDE Clusters 31, 38, 34, 46, 12, and 22 were then projected onto the cell-level UMAP (**Supplementary Figure 5B**). Each group of perturbations represented by a cluster is observed to colocalize in a well-defined region of the UMAP, distinct from the NTC cells (**Figure 2C**), indicating a potential change in cell state. Adjacent clusters in the MDE additionally colocalize in the UMAP. Notably, the clusters associated with pluripotency and developmental processes (Clusters 31 and 38) are positioned distinctly from the NTC on the UMAP, suggesting a shift in cell state away from the pluripotent state represented by the NTC cells. As the UMAP is based on highly variable genes, and not specifically tailored to DEGs, this observation corroborates the significant impact of these perturbations on altering the cellular identity as identified in **Figure 1F**. We then overlaid the median number of DEGs per perturbation for each cluster of the cluster-level MDE. The clusters inducing the strongest transcriptional change are Clusters 31 (Pluripotency Factors), 38 (Development), and 49 (Pluripotency Factors) (**Supplementary Figure 5C**). Finally, the bulk fitness screen (**Figure 3**) was leveraged to assess relationships between MDE functional clusters and fitness impact. Over time, a progressive negative fitness impact was observed in clusters with mitochondrial function (8, 42, 20, 46, and 34), the BAF complex cluster (Cluster 40), and the MTOR and endosome cluster (Clusters 46, 34) (**Supplementary 5D, 5E**). This suggests that perturbations in these clusters increasingly disrupt cell fitness, particularly affecting processes linked to mitochondrial function, chromatin remodeling, highlighting their critical roles in maintaining the cellular fitness of hiPSCs.

### A ML Architecture Towards Rationale Cell State Engineering of hiPSCs

Recently, there have been a number of ML models developed that aim to *in silico* predict the effect of a perturbation on cell state, such as GEARS^71^ and CellOT^72^, among others. Generally speaking, these models aim to learn a representation of the perturbation applied (a graph embedding in the case of GEARS, an optimal transport map in the case of CellOT) to predict the resultant cell state. Inspired by these models, we wondered if in the specific context of hiPSCs, a cell line primed for cell state engineering, there was a unique opportunity to flip this paradigm. That is, can a model predict the perturbation(s) that need to be applied to the wild type (WT) cell to achieve a given cell state of interest?

Towards this goal, we propose a general model architecture that learns specifically from perturbation atlases (**Figure 4C left**). During the training phase, the perturbation atlas is split into perturbed and unperturbed (NTC) cells, which are then passed into a self- or semi-supervised (e.g., gene ontology graph-informed) cell-state encoder to learn a representation (low-dimensional vector embedding) of the perturbed cell state – that is, what makes this perturbed cell state different from the control cells? The encoder is not informed about the identity of the perturbation made to the perturbed cell. This embedding is then passed into a supervised perturbation decoder, which is trained using the ground-truth labels from the perturbation atlas, to determine the perturbation(s) with the highest probability of pushing the WT cell towards this perturbed cell state. Then, for inference (**Figure 4C right**), the transcriptome of desired cell state (e.g. bulk-RNA seq of a neuronal population) can be passed through the encoder-decoder architecture to predict perturbation(s) with high log-probabilities of inducing that cell state. These perturbations can be experimentally validated, and results can be used to improve model performance. We coin this architecture CHEF (<u>C</u>ontrastive <u>H</u>uman-cell <u>E</u>ngineering <u>F</u>ramework), as it predicts perturbation recipes driving a desired cell state.

As a proof of principle implementation of this architecture, we used ContrastiveVI^73^ as the cell-state encoder, and a simple logistic regression classifier as the perturbation decoder (**Figure 4C middle**). ContrastiveVI is a variational autoencoder (VAE) framework that learns a representation specific to perturbed cells that isolates them from control cells called the “salient embedding.” Given its unsupervised nature, the model is inherently generalizable to learning cell state representations of unseen transcriptomes for which the associated perturbation is unknown. Further, an important feature of ContrastiveVI is that as input it takes raw, unnormalized RNA-seq profiles. This improves the utility of this model far beyond simply using the learned MDE to co-cluster a cellular profile near similar perturbations, as the latter requires the same normalization, processing, and batch-correction as the data its trained on (i.e., unseen data would necessarily need to be from the same experiment). Representations learned from this model are then passed into a supervised logistic regression classifier, which is trained to predict the associated perturbation.

The training loop and parameters we used are outlined in **Supplementary Figure 4F**. We initially subset our data down to 33 strong perturbations with at least 100 DEGs and 100 associated cells as our training set to ensure the model was trained on perturbations with a strong phenotypic effect and high sample representation within the dataset. Class balancing was performed by subsampling to 100 cells per perturbation, and feature selection was performed by using the same top 2,000 highly variable genes used to create the MDE. Raw (unnormalized) counts were then passed into the ContrastiveVI model with the batch as a categorical covariate. Hyperparameters were optimized using RayTune/Hyperband^74^ and the model was finally trained until the validation loss (negative ELBO) no longer improved. The salient embeddings from the model were visualized using UMAP (**Figure 4D**), where we observed that the ContrastiveVI model was able to generate similar embeddings for NTC cells that separated them away from perturbed cells. Then, a logistic regression classifier was trained on the salient embeddings to predict the associated perturbation with a 80/20 train-test split, with the confusion matrix and performance statistics reported in **Figure 4E**. The model exhibited moderate classification accuracy of 25%, which increased to 39% when considering the top 3 accuracy. In our view, this is rather respectable performance as this represents perturbation classification of the transcriptional profile of a single cell. At random, the classification probability would be 1/34 or ∼3%. Further, we noted that related perturbations were being confused for each other; for example, JOSD1 is confused for POU5F1, which co-cluster in the MDE.

As the ContrastiveVI model contains a regularization term to ensure the salient embedding is represented by a multivariate normal gaussian with identity covariance and the loss function is evaluated by the KL-divergence between embedding spaces (in this special case, the euclidean distance between their means), distances in salient embedding space are representative of distances in the input space. We therefore computed the average salient embedding of each training perturbation to represent the salient embedding of the pseudo bulk profile of cells with the same perturbation. A logistic regression model was then fit to these pseudo bulk profiles to serve as a pseudo bulk perturbation decoder. To simulate previously unseen, pre-validated, bulk-RNA seq profiles of desired transcriptional states, we used pseudo bulk transcriptional profiles of the remaining strong perturbations not previously seen by the model. These perturbations were passed into the pretrained ContrastiveVI encoder, and pseudo bulk perturbation decoder. Classification probabilities for unseen perturbations with class probability for any one class >10% were visualized in a heatmap (**Figure 4F**). We observed that for every unseen perturbation visualized, the model suggested perturbations from either the same or adjacent clusters in the MDE, validating the model predictions. The predictions spanned eight different MDE clusters. Further, we observed that unseen perturbations classified as POU5F1 (such as PTPN1) were additionally classified as JOSD1 (same MDE cluster), and FLYWCH2 (adjacent MDE cluster).

We additionally created a “naive” model to demonstrate the utility of the current implementation of CHEF. Put simply, the naive model represents a case where one would simply “look up” perturbations with similar DEGs to their desired cell state, rather than learning an embedding of the cell state. In the naive model, we used as features the union of all DEGs of all strong perturbations (5665 unique DEGs). Each perturbation was represented by a binary vector of these perturbations, with a one in indices of DEGs of that perturbation, and zeros otherwise. NTC was represented by a vector of zeros. We then fit a logistic regression classifier on the same training perturbations and used to evaluate the same unseen perturbations as in **Figure 4F**. We observed that the naive model classified most unseen perturbations as NTC, demonstrating the advantage of learning cell state embeddings (see Supplementary Data). Overall, our CHEF implementation represents a proof of principle of an inference architecture for cell state engineering of hiPSCs.

### Comparison of Protein-Protein Interactions in hiPSCs Using Size-Exclusion Chromatography-Mass Spectrometry (SEC-MS) and Perturb-Seq

Perturb-Seq analysis in iPSCs enabled us to classify perturbations into clusters based on their transcriptomic profiles. This approach revealed strong transcriptomic associations among several protein complexes, including mitochondrial ribosome subunits as well as the Ragulator complex. Notably, clusters corresponding to mitochondrial translation functions show high transcriptomic correlation within each cluster, and our data suggest that perturbations particularly to the MDE functional clusters of mitochondrial function (Clusters 8, 42, 20, 46, and 34), the BAF complex (Cluster 40), and the MTOR and endosomes (Clusters 46, 34) (see above; **Supplementary 5D, 5E**) increasingly disrupt cell fitness. To validate that these transcriptomic clusters represent actual protein complexes in iPSCs, we conducted size-exclusion chromatography coupled with mass spectrometry (SEC-MS) in KOLF2.1J iPSCs. SEC-MS separates protein complexes by molecular weight and uses a guilt-by-association framework to assign co-eluting proteins into the same complex^75,76^ (**Figure 5A**). The SEC-MS data were then analyzed for cosine similarity to assess the systematic co-fractionation of proteins across fractions as physical evidence of their interactions (see Methods for details). We found significant enrichment in high cosine similarities between pairs of proteins within clusters of mitochondrial function and MTOR (Clusters 34, 41, 42, 46 and 48) from our Perturb-Seq data with the cosine similarity heatmap showing distinct clustering patterns (**Figure 5B**) with strong internal similarity within the previously identified clusters from the Perturb-Seq MDE. For example, proteins corresponding to the transcripts in clusters related to mitochondria, e.g. Cluster 46 (MT Complex 1) and Clusters 42 and 41 (Mitochondrial ribosomal L proteins; MRPL), co-eluted across multiple fractions, confirming the close physical association of mitochondrial ribosome subunits. The mTORC signaling Cluster 34, encompassing the Ragulator-Rag complex which consists of LAMTOR subunits and regulates MAPK and mTORC1^77,78^ displayed high cosine similarity among LAMTOR subunits and the fractionation profiles of LAMTOR2, LAMTOR3, LAMTOR4, and LAMTOR5 (**Figure 5E**) showed overlapping elution peaks, confirming their co-existence in the same molecular weight fractions and indicating a stable protein complex in iPSCs. Further visualization of these clusters as PPI networks (**Figure 5C**) highlights the interactions within a cluster, e.g. Ragulator components in Cluster 34, as well as among clusters, with strong connectivity observed among MRPL subunits (Clusters 42 and 41), mitochondrial ribosomal small (MRPS) subunits (Cluster 48). Clusters 42, 41, and 48, positioned closely on the Perturb-Seq MDE (**Supplementary Figure 5A)**, highlighting this interconnectivity at a functional level and reflected by the strong transcriptional coherence in the co-expression patterns within these perturbation clusters (**Figure 5D)**. Interestingly, Cluster 34 shows particularly tight correlation of gene expression among Ragulator components, indicating that knocking down any of the subunits leads to the same phenotype, suggesting overall perturbation of the Ragulator complex function. Overall, the SEC-MS data validated the PPIs inferred from Perturb-Seq, such as Ragulator complex, mitochondrial ribosome and MT complex 1, confirming that the observed transcriptomic clusters can correspond to physically stable protein complexes in iPSCs. This integrative analysis demonstrates that transcriptomic clustering from Perturb-Seq can provide insights into protein complex formation such as the functional organization of mitochondrial and signaling-related complexes in stem cells while also providing valuable information of multiple clusters of functionally related proteins without evident or previously known physical association.

**Figure 5.**
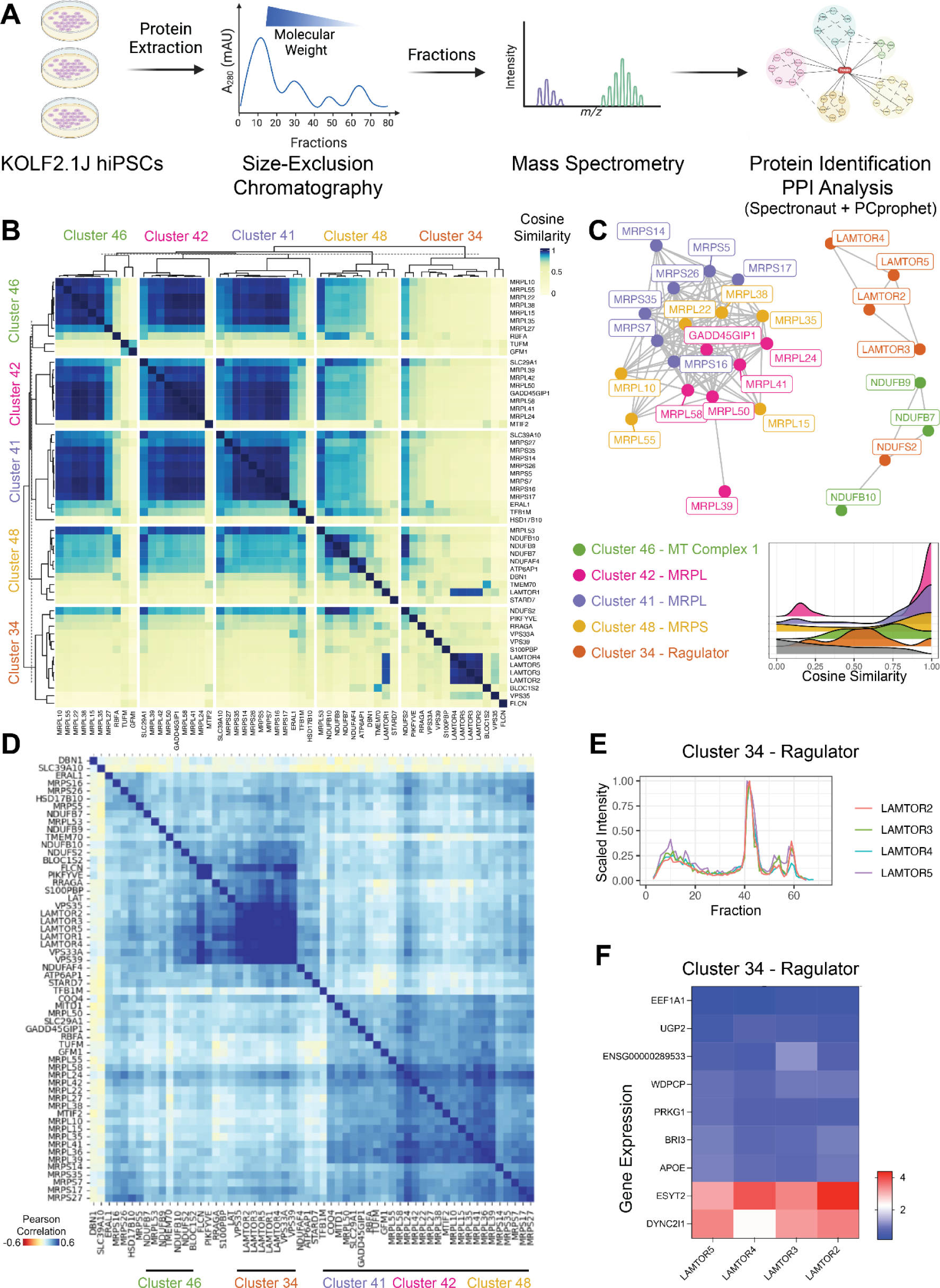
Validation of Protein-Protein Interactions in hiPSCs Using Size-Exclusion Chromatography-Mass Spectrometry (SEC-MS) and Transcriptomic Analyses. **(A)** Experimental workflow for protein interaction analysis in KOLF2.1J hiPSCs. Proteins were extracted from hiPSC samples and separated by size-exclusion chromatography. Fractions were analyzed by mass spectrometry to identify and quantify proteins by Spectronaut, followed by protein-protein interaction (PPI) analysis yielding interaction networks based on cosine similarity of fractionation profiles. **(B)** Heatmap of cosine similarity among Perturb-Seq derived MDE protein clusters that show significant enrichment for very high cosine similarity in the SEC-MS PPI. Perturb-Seq derived MDE Cluster 46 (green), Cluster 42 (pink), Cluster 41 (blue), Cluster 48 (purple), and Cluster 34 (orange). Cosine similarity scores are shown, with higher similarity (yellow) indicating co-elution across fractions. **(C)** Identified interactions among proteins from SEC-MS, with each color-coded according to Perturb-Seq MDE cluster. Clusters are labeled as follows: Cluster 46 (MT Complex 1, green), Cluster 42 (MRPL, pink), Cluster 41 (MRPL, blue), Cluster 48 (MRPS, purple), and Cluster 34 (Ragulator, orange). Edges represent strong cosine similarity (>0.97), indicating probable protein interactions. Histogram graph showing cosine similarity distribution of protein pairs within individual clusters (color-filled) versus all protein pairs between any different Perturb-Seq MDE clusters (grey filled), with peaks near 1.0 indicating strong interaction scores within clusters. **(D)** Pearson correlation heatmap of transcriptomic data from Perturb-Seq across MDE derived cluster: *Cluster* 46 (green), Cluster 42 (pink), Cluster 41 (blue), Cluster 48 (purple), and Cluster 34 (orange). **(E)** Fractionation profile of Ragulator complex subunits (Cluster 34), which create a connected network component in **C**, across size-exclusion chromatography fractions. **(F)** Gene expression heatmap Log2FoldChange of common DEGs between LAMTOR2, LAMTOR3, LAMTOR4, and LAMTOR5.

### Identification of New Genes Involved in Mitochondrial Metabolism

The Perturb-Seq atlas serves as a powerful resource not only for identifying protein complexes, but also for uncovering new gene function on a broader scale. The pairwise Pearson correlation heatmap of pseudo-bulk transcriptomes for each perturbation reveals distinct clusters where known protein complexes co-cluster together, demonstrating correlated gene expression patterns among subunits (**Figure 6A**). Two prominent clusters emerged from these analyses: one enriched with key pluripotency factors (Cluster 31), and another characterized by genes involved in mitochondrial translation (Cluster 20). Within the mitochondrial translation cluster, we identified several novel genes, such as ZBTB41, PRR13, UBFD1, LFNG, and BOD1L1 which show high correlation with known mitochondrial factors, suggesting potential roles in mitochondrial metabolism. We focused on ZBTB41 as it is a poorly characterized zinc-finger transcription factor, and generated KOLF2.1J cell lines transduced with sgRNA NTC and targeting MRPL11, MRPL37, or ZBTB41. The knockdown of MRPL11, MRPL37, and ZBTB41 was first validated through qPCR, confirming significant repression of target gene expression relative to the control (sg-NTC) (**Supplementary Figure 6B**). All perturbations showed robust reduction in their respective mRNA levels, demonstrating the efficacy of the knockdowns. Next, we explored the genomic signature of these perturbations by analyzing gene expression patterns associated with mitochondrial function (**Supplementary Figure 6C**). Perturb-Seq data revealed downregulation of key mitochondrial genes (e.g., MT-ND2, MT-ND1) upon disruption of MRPL11, MRPL37, and ZBTB41, further indicating these perturbations are linked to mitochondrial dysregulation; MT-ND1 and MT-ND2 upregulation under MRPL11, MRPL37, and ZBTB41 knockdown was also confirmed (**Supplementary Figure 6G, 6H**).

**Figure 6.**
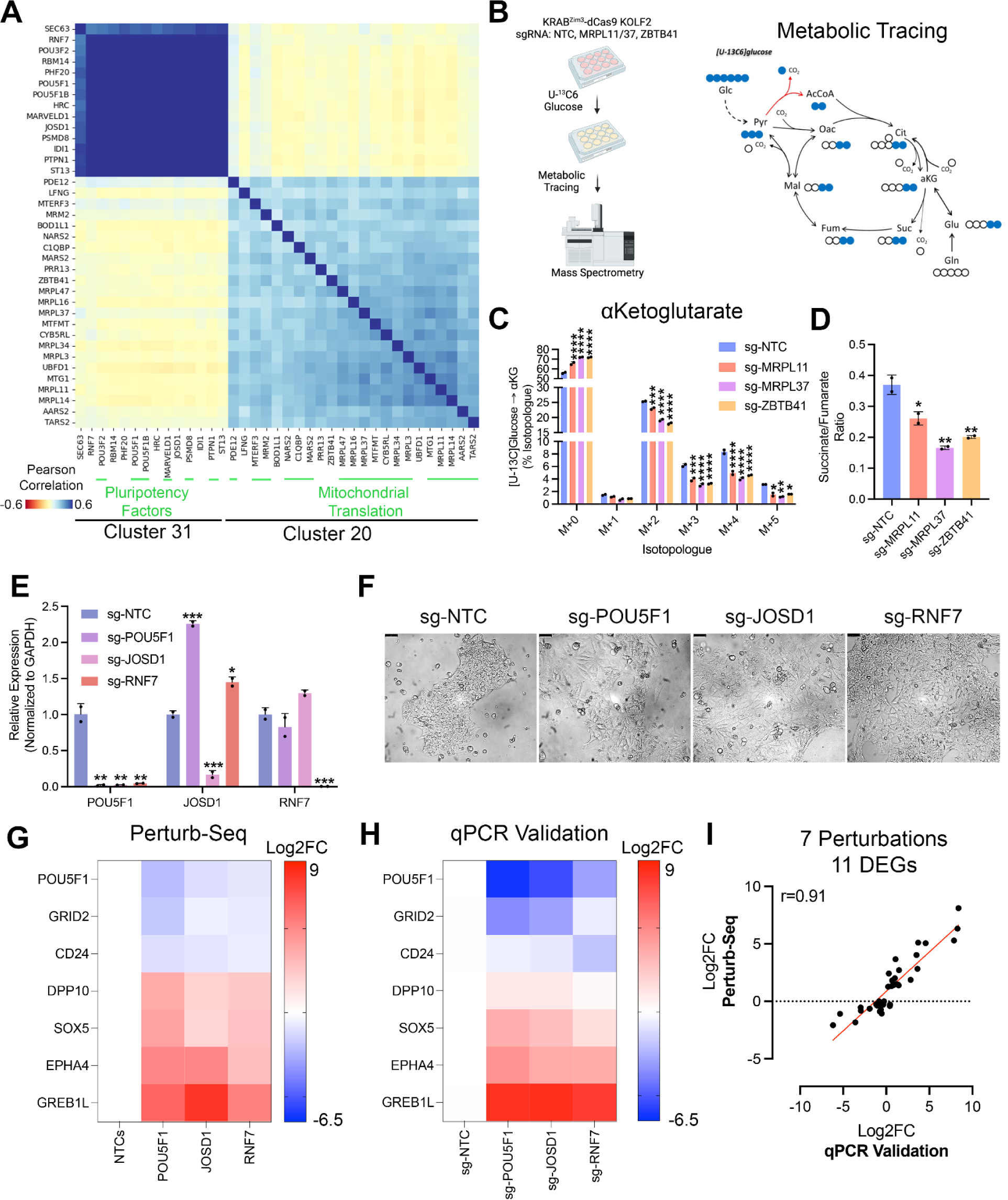
Validation of Novel hiPSC Pluripotency and Mitochondrial Factors. **(A)** Pearson correlation between perturbation MDE clusters with known interactors highlighted in green for cluster 31/Pluripotency factors and cluster 20/Mitochondrial Translation. **(B)** Experimental workflow for metabolic tracing of mitochondrial flux validation experiment run in KOLF2.1J KRAB^ZIM3^-dCas9 transduced with sgRNAs NTC or MRPL11, or MRPL37 or ZBTB41 **(C)** Metabolic tracing of [U-13C]Glucose into α-Ketoglutarate isotopologues (M+0 to M+5) under different genetic perturbations. Five different single-guide RNAs (sgRNAs) were used to target MRPL11 (orange), MRPL37 (purple), ZBTB41 (yellow), and a non-targeting control (NTC, blue). The bar graph represents the percentage of isotopologue labeling from [U-13C]Glucose, showing the distribution from M+0 (unlabeled) to M+5 (fully labeled) α-Ketoglutarate. Significant differences are indicated between sgRNA-targeted genes and the NTC, as denoted by the asterisks (*p < 0.05,**p < 0.01,***p < 0.001, ****p < 0.0001). **(D)** Ratio of succinate to fumarate in cells transduced with various sgRNAs: MRPL11 (red), MRPL14 (green), MRPL37 (purple), ZBTB41 (yellow), and a non-targeting control (NTC, blue). Significant differences are indicated between sgRNA-targeted genes and the NTC, as denoted by the asterisks (*p < 0.05, **p < 0.01). **(E)** *Quantitative PCR analysis of gene expression* in cells transduced with sgRNAs targeting POU5F1 (purple), JOSD1 (pink), RNF7 (red), and a non-targeting control (NTC, blue). Asterisks indicate statistical significance (*p < 0.05, **p < 0.01, ***p < 0.001). **(F)** KOLF2.1J KRAB^ZIM3^-dCas9 were transduced with NTC, POU5F1 or JOSD1 sgRNAs and imaged 4 days post-transduction with brightfield at 20x magnification. Top left black scale bar = 50 µm. **(G)** Log2 fold change (Log2FC) in gene expression from Perturb-Seq across different sgRNA treatments. The heatmap displays the Log2FC of genes (POU5F1, GRID2, CD24, DPP10, SOX5, EPHA4, GREB1L) in cells treated with sgRNAs targeting POU5F1, JOSD1, and RNF7 compared to non-targeting controls (NTCs). **(H)** Log2 fold change (Log2FC) in gene expression from qPCR validation across different sgRNA treatments. The heatmap shows the Log2FC of the same genes as in panel G, validated by qPCR in cells treated with sgRNAs targeting POU5F1, JOSD1, and RNF7 compared to sg-NTC. **(I)** Correlation between Log2FC values from Perturb-Seq and qPCR validation. The scatter plot shows the correlation of Log2FC values from Perturb-Seq and qPCR validation, with a Pearson correlation coefficient (r = 0.91) across 7 perturbations and 11 differentially expressed genes (DEGs).

We further explored the role of these genes by performing a metabolic tracing experiment (**Figure 6B**). Cells were treated with [U-^13^C_6_]Glucose, and the incorporation of labeled glucose into α-Ketoglutarate (αKG) was measured using gas chromatography-mass spectrometry. Perturbations in MRPL11, and MRPL37, as well as ZBTB41, resulted in significant alterations in the distribution of αKG isotopologues (**Figure 6C**), including an increase in M+0 that indicates a disruption in tricarboxylic acid (TCA) cycle flux. This finding was further supported by a similar trend in isotopologue labeling for other key TCA cycle intermediates, including citrate, fumarate, and malate (**Supplementary Figures 6D, 6E, 6F**). In addition, we also examined the succinate/fumarate ratio as another indicator of mitochondrial function (**Figure 6D**). Compared to sg-NTC, cells perturbed for MRPL11, MRPL37, and ZBTB41 displayed a significant reduction in the succinate/fumarate ratio, further supporting the hypothesis that these perturbations affect mitochondrial metabolic processes. The labeling patterns of the TCA cycle metabolites suggest impaired glucose oxidation and mitochondrial function across these perturbations. Overall, this integrated analysis of gene expression, metabolic flux provides strong evidence that perturbations of MRPL11, MRPL37, and ZBTB41 disrupt mitochondrial metabolism, as reflected by both the gene expression changes in mitochondrial genes and the altered metabolic flux through the TCA cycle. This analysis highlights the role of previously uncharacterized genes like ZBTB41 in mitochondrial function and underscores the broader connection between mitochondrial ribosomal protein function and metabolic regulation.

### Identification of New Genes Involved in Pluripotency Maintenance

Among the 50 clusters identified from the MDE analysis, Cluster 31 stood out with 14 perturbations, many of which target genes with known roles in pluripotency maintenance or differentiation regulation (**Figure 6A**). This cluster includes essential pluripotency factors, such as POU5F1, which is crucial for maintaining the undifferentiated state of stem cells. Additionally, Cluster 31 includes PTPN1 and POU3F2 (also known as BRN2), which have been implicated in neuronal differentiation^31,79^, suggesting a potential shift towards a neural lineage when perturbed. Other perturbations within Cluster 31, including JOSD1, RNF7, RBM14, PHF20, and ST13, have not previously been linked to pluripotency or differentiation, making their association with known pluripotency regulators intriguing. This clustering suggests these genes may participate in novel pathways governing stem cell maintenance or lineage specification. Given the co-clustering of JOSD1 and RNF7 with well-known pluripotency factors, we performed JOSD1 and RNF7 knockdown experiments in the KRAB^ZIM3^-dCas9 KOLF2.1Js to test their involvement in pluripotency maintenance with sgRNA targeting POU5F1 (sg-POU5F1) as a positive control and NTC gRNA (sg-NTC) as a negative control. We first assessed knockdown efficiency by qPCR and observed robust repression for all target genes (**Figure 6E**). Interestingly, we found that POU5F1 expression was reduced not only in sg-POU5F1 cells but also in cells where JOSD1 and RNF7 were repressed. However, knockdown of POU5F1 did not lead to reduced expression of JOSD1 or RNF7. Morphologically, we observed notable changes in the sg-POU5F1, sg-JOSD1 and sg-RNF7 cells compared to controls sg-NTC. These cells exhibited a loss of typical stem cell colony structure, showing a more spread-out and flattened appearance, with larger individual cell size (**Figure 6F**). This morphology is characteristic of cells undergoing differentiation, suggesting that repression of JOSD1, RNF7 and POU5F1 disrupts the pluripotent state and drives the cells towards a differentiated phenotype. To explore the transcriptional impact of these knockdowns, we analyzed differentially expressed common DEGs from Perturb-Seq found in Cluster 31. The DEG profile for Cluster 31 perturbations, including JOSD1 and RNF7, showed a decrease in key pluripotency markers such as POU5F1, GRID2, and CD24, alongside an increase in differentiation markers including SOX5, EPHA4, and GREB1L (**Figure 6G**). These transcriptional changes, indicative of a shift from a pluripotent state towards differentiation, were validated in our single-plex validation samples, confirming that JOSD1 and RNF7 knockdown drives a similar transcriptional response to that of POU5F1 knockdown (**Figure 6H**). Interestingly, while both JOSD1 and RNF7 knockdowns led to substantial transcriptional and morphological changes associated with loss of pluripotency, the fitness screen did not reveal any significant impact on cell viability for POU5F1, JOSD1, or RNF7 knockdowns (**Supplementary Figure 6I**). This highlights the specificity of these genes in maintaining pluripotency rather than cell survival, and it underscores the unique value of Perturb-Seq in identifying these novel pluripotency regulators that may not impact fitness directly. Finally, a strong correlation (r=0.91) was observed between the Perturb-Seq screening data and low-throughput qPCR validation (**Figure 6I**), underscoring the accuracy and reliability of Perturb-Seq in capturing gene expression changes. This robust correlation validates Perturb-Seq as an effective approach for identifying novel pluripotency regulators, such as JOSD1 and RNF7, within complex transcriptional networks in hiPSCs.

## DISCUSSION

In this study, we performed a comprehensive genome-wide CRISPRi Perturb-Seq screen in KOLF2.1J iPSCs, targeting 11,739 genes in parallel to systematically dissect the transcriptional and functional landscape of pluripotent stem cells. Using strong perturbations and transcriptomic profiling correlations projected via MDE, we were able to recapitulate established protein complexes and identify genes with redundant functions, providing a detailed map of functional relationships within iPSCs (**Figure 2**). Our approach identified key regulators of pluripotency and revealed how perturbations influence cellular fitness (**Figure 3**) and transcriptional signatures (**Figure 6**). SEC-MS in KOLF2.1J iPSCs validated PPIs identified from the Perturb-Seq derived MDE **(Figure 5)**. We further leveraged our Perturb-Seq dataset to train a machine learning model that predicts cell states of unseen perturbations from the training dataset (**Figure 4**), demonstrating a potential cell state engineering application of the perturbation atlas. Notably, the strongest hits from the screen included well-established pluripotency factors such as POU5F1 and NANOG, validating the screen’s ability to capture critical regulators. Additionally, we discovered new metabolic factors that co-clustered with mitochondrial gene perturbations and identified a cluster of pluripotency factors with new perturbations inducing phenotypes similar to POU5F1 repression (**Figure 6**).

Through this Perturb-Seq dataset in iPSCs, we were also able to identify new genes involved in pluripotency maintenance. In particular, our MDE analysis identified Cluster 31, which contains well-known regulators of pluripotency, such as POU5F1, and novel candidates like JOSD1 and RNF7, both of which showed similar effects to POU5F1 knockdown. This finding suggests that JOSD1 and RNF7 play a previously unrecognized role in pluripotency maintenance, supported by both morphological changes and transcriptional shifts indicative of differentiation upon its knockdown. JOSD1, a deubiquitinase, and RNF7, a catalytic subunit of an E3 ligase complex, are pivotal for protein homeostasis by modulating ubiquitination dynamics, with JOSD1 reversing ubiquitin modifications^80^ and RNF7 facilitating neddylation and substrate recruitment for degradation^81,82^. This functional interplay suggests that disrupting these ubiquitin-related pathways could destabilize the tightly regulated protein turnover critical for maintaining pluripotency, as protein quality control is essential in stem cells to preserve a balanced proteome and stem cell identity^83^. Despite their influence on pluripotency, our fitness screen (**Supplementary Figure 6I**) did not indicate that POU5F1, JOSD1 and RNF7 significantly impact overall cell fitness. Our comprehensive fitness profiling revealed that while many genes are essential for pluripotency, others play distinct roles in maintaining cell fitness. For example, genes involved in chromatin remodeling (e.g., SAGA complex members) were enriched in our fitness screens, indicating that their perturbation enhances cell proliferation while potentially compromising pluripotency. In particular, TADA2B, a core member of the SAGA complex, is the top1 enriched hit from our fitness screen in KOLF2.1Js (**Figure 3**). Recent literature has shown that TADA2B-associated chromatin changes influence pluripotency and differentiation independently, with perturbations in the SAGA complex components favoring cell fitness without necessarily supporting pluripotent identity^57,63^. This reinforces the idea that fitness and pluripotent identity are governed by different pathways and cannot be defined by cell fitness alone. We also observed that the Fanconi anemia pathway and mitochondrial translation genes were critical for iPSC survival, underscoring the heightened sensitivity of iPSCs to metabolic and DNA repair disruptions. Our findings align with the concept that pluripotency dissolution involves a “push-pull” mechanism^63^, where the loss of pluripotent identity occurs independently of differentiation signaling. This is consistent with our identification of chromatin regulators as key players in pluripotency dissolution, while developmental transcription factors were more strongly associated with differentiation signaling.

We also acknowledge certain limitations of this study. As previously discussed, the Perturb-Seq assay is inherently biased towards capturing perturbations that do not induce a negative fitness impact. To have sufficient coverage / statistical confidence, we filter for cells that, in the best case scenario, have 25 associated cells pre-filtering. As a result of our QC/filtering pipeline, most perturbations require more than 25 starting cells to pass this threshold, which is challenging to achieve for perturbations with deeply negative fitness impacts (**Figure 3G**). Hence, this study may not fully capture the transcriptional effects of some key pluripotency drivers. This is further influenced by the choice of pluripotency maintenance media containing self-renewal factors such as hFGF and hTGFβ. Inherently, the media conditions are promoting maintenance of the stem cell in a pluripotent state, and thus any perturbations have to overcome the self-renewal effect of the media environment. As all perturbations are assayed in a multiplexed fashion, a key confounding factor diminishing the resolution of characterizing individual perturbations may be external stimuli from cell-cell signaling.

Additionally, while our experimental and analysis pipelines do recapitulate well characterized biological processes (**Figure 2B**), driving confidence that we capture some aspects of true biological signal, we feel there is certainly room for improvement. First, experimentally we have a median 5000 UMIs per cell. While our sequencing depth is above the recommended 1000 UMIs per cell from Peidli et al.^18^, achieving higher sequencing depths closer to ∼10,000 UMIs/cell^15^ will enable finer grain resolution. Second, in our single-plex validations through qPCR (**Supplementary Figure 4F**), we confirmed robust repression of target genes via the KRAB^ZIM3^-dCas9 system, and therefore anticipate that we achieve sufficient perturbation magnitude. However, a dual sgRNA library^15^ could be utilized in the future to ensure stronger perturbation magnitude. Additionally, there is a possibility of confounding effects from CRISPRi off-targets affecting neighboring genes, enhancers, insulators, and potentially pseudogenes as well as non-coding RNAs.

We seek to emphasize the importance of selecting a substantially large proportion of the screen to comprise of NTC sgRNA, and of having the same NTC sgRNA present in each batch. In our screen, NTC sgRNA comprised ∼4% of the library, and were shared across all 3 batches. We observed in our fitness data that though most NTC sgRNA are benign, some aberrant sgRNA can have drastic negative fitness effects (as much as −2.5 Z, **Figure 3F**). Additionally there may be single cells that receive a benign NTC sgRNA, though nevertheless display a deviant phenotype (**Supplementary Figure 1D**). We envision hiPSCs being particularly prone to this phenomenon as they are capable of spontaneous state change / differentiation, and hypothesize that a population of cells in the pluripotent state have more cell-to-cell variability than in the terminally differentiated state. As a result of these factors our starting population of ∼150K NTC cells was reduced to ∼30K NTC cells post filtering (∼10K cells / batch), emphasizing the importance of ensuring a sufficiently large population of NTC cells. We recommend that for future screens, in an ideal case at least 5% of the sgRNA library should consist of NTC sgRNA. When feasible, experiments should be performed in a single batch (i.e. 10X run) to avoid the potentially strong confounding effect of batch effect (**Supplementary Figure 1H**). In cases where this is not experimentally feasible (i.e. at the genome scale), we found, as others^15^ that the strategy of leveraging batch level internal controls (i.e. having the same set of NTC sgRNA in each batch) is effective at reducing batch effects (**Supplementary Figure 1H**).

Another key bottleneck for this study (and we believe fundamentally all single-cell studies) is the ability to generate faithful cell state embeddings. While we chose the top 2000 highly variable genes as features to get a global picture of phenotypic variation, it is unclear whether this is the optimal feature set to separate perturbation-specific effects. Additionally, we observed that using the top 50 principal components (PCs) as features for each cell, which is the norm in the single-cell field, only accounted for < 10% of the total variation in the dataset. We hypothesize that this is a result of Perturb-Seq focusing on a single cell line, which is much more homogenous than the heterogeneous cell type mixture in (for example) PBMCs, where the top 50 PCs perform well. These two factors undoubtedly diminished our ability to resolve the fine-grain effect of perturbations on phenotype. In our opinion, learning robust cell embeddings is an open problem of seminal importance to the Perturb-Seq field.

As outlined by Peidli et al.^18^, there is to date no standardized Perturb-Seq analysis pipeline, and given the >11,000 perturbations we assayed, we sought to develop a method that is scalable and intuitive. Sources of heterogeneity emerged at both the sgRNA and cell level. Despite the modest cell state representations learned by PCA, we still observed heterogeneity in phenotypic profiles induced by individual perturbations. This was a key challenge that we aimed to address with our computational pipeline. We noted instances where for a given perturbation, while all three sgRNA did induce sufficient target knockdown, each sgRNA would induce phenotypic changes of variable magnitudes. As a part of our pipeline, we leveraged the energy distance to identify sgRNA likely inducing phenotypic alterations. One limitation at the genome scale is the computational complexity of running the energy distance test (i.e. permutation test with 10,000 permutations per sgRNA) for all >30,000 perturbing sgRNA. To engineer around this we decided to use a bootstrapping approach to generate a null distribution from 10,000 random samples of NTC cells to identify an upper-quartile cutoff of energy distance. sgRNA with energy distances exceeding this cutoff were admitted. This reduced the number of energy distances to be computed from on the order of ∼300 million to ∼50 thousand. Further, cells experiencing incomplete perturbation penetrance are challenging to identify. As one example, cells may appear artificially repressed for their target gene as an artifact of a scRNA-Seq dropout event, and inclusion of these cells can mask perturbation-specific effects. The use of CRISPRi adds an additional consideration, as cells may experience repression of variable magnitude. While Mixscape ^17^ is in our opinion a valuable tool for identifying cells escaping perturbation with high resolution, we found that at the genome scale it was challenging to execute with reasonable runtime. Hence, we developed a Mixscape-inspired method where we first fit a One-Class SVM to NTC cells (just as Mixscape fits a Gaussian to NTC perturbation scores) and used it to predict the probability of a perturbed cell being in the NTC population, using the same threshold as at the sgRNA level to filter. While there is certainly room for improvement, we feel that our approach helped to resolve perturbation specific effects. After filtering for perturbations with at least 25 remaining cells and >10 DEGs, we observed that the identified 1332 strong perturbations did not co-cluster highly with NTC cells on the UMAP (**Figure 2D**), noting that the UMAP is computed on 2000 highly variable genes (not perturbation specific DEGs) as features.

The demonstrated ML model, in its current implementation, is quite limited in its capabilities. We subsetted down to training perturbations with a high number of samples and large phenotypic impact to achieve fair model performance. We found that including perturbations with fewer total cells tended to lead to overfitting, and overall poor inference. As a result, the model can only suggest between 34 total perturbations to make. We anticipate that several improvements can be made, such as bootstrapping to generate more synthetic data points, greater fine-tuning of model hyperparameters, and use of semi-supervised learning in the encoder to achieve more robust inference for a greater number of perturbations. Further, this model is not capable of predicting combinations of perturbations, but rather individual perturbations that may be assayed in combinations. Nevertheless, we argue that this model still demonstrates an important initial result, as it is capable of accurately suggesting perturbations across a diversity of cell states. To our knowledge, this encapsulates the first proof of concept model capable of predicting perturbations driving a cell state, and we hope the broader scientific community can iterate and improve on this basal architecture. Future directions may include integration of the hiPSC CRISPRi dataset (this study) with the hESC mORF overexpression^6^ dataset to encapsulate genetic perturbations in both directions, and use of graph neural networks to incorporate external knowledge from biological knowledge graphs (such as protein-protein interaction networks) to enhance the model towards predicting combinatorial perturbation recipes. A model of this capability, as previously outlined^84^, would be invaluable for deterministic cell state engineering.

Overall, our study represents a comprehensive perturbation cell atlas of hiPSCs. We feel that this dataset is a rich resource for the scientific community to query the complexities of pluripotency. As an example, we leveraged a MDE map of correlated perturbation profiles to identify novel pluripotency factors. Future studies could include reanalysis of the dataset to focus on genes relevant for a differentiation trajectory of choice, and investigating how perturbations push the undifferentiated stem cell towards that trajectory. For example, combining scRNA-Seq in human-derived teratoma model^85^ with perturbations would be an efficient method to assess and identify important genes involved in hPSCs differentiation. Further, we envision this dataset to be expanded into a multiomic atlas, integrating data from (for example) chromatin structure^86^ and spatial^87^ information. Finally, we anticipate great utility in combining datasets across perturbation cell atlases^88^ to form a comprehensive human cell perturbation atlas to unify human cell biology.^19^ We hope that this resource and the associated computational methods developed is used by the broader scientific community to unravel previously unknown biology of the pluripotent stem cell.

## ACKNOWLEDGEMENTS

The authors would like to thank Adithya Vellal, Michael Tong, and Joseph Rainaldi for useful discussions and support throughout all stages of the study. This work was generously supported by the NIH Bridge2AI program (OT2OD032742) as well as other NIH grants (R01HG012351, R01NS131560, U54CA274502), a Department of Defense Grant (W81XWH-22-1-0401), CIRM training grant (EDUC4-12804), UCSD Institutional Funds, and a Rubicon grant from Dutch Research Council (NWO, 019.231EN.013). This publication includes data generated at the UC San Diego IGM Genomics Center utilizing an Illumina NovaSeq 6000 that was purchased with funding from a National Institutes of Health SIG grant (#S10 OD026929) National Cancer Institute (P30CA023100). Parts of some figures were illustrated with BioRender.

## AUTHOR CONTRIBUTIONS

Conceptualization and Design: SN, YD, PM; Experiments: SN, YD, AD, AF, YHL, BC, MM, HK, EP, YZ, AS, JNH; Computational analyses: YD, SN, BP, JG; Supervision: JM, KO, TC, JYC, CM, EL, TI, NK, PM; Writing: SN, YD, PM with input from all authors.

## DATA AVAILABILITY

Code and raw data are available via P.M.

---

**Supplementary Figure 1:**
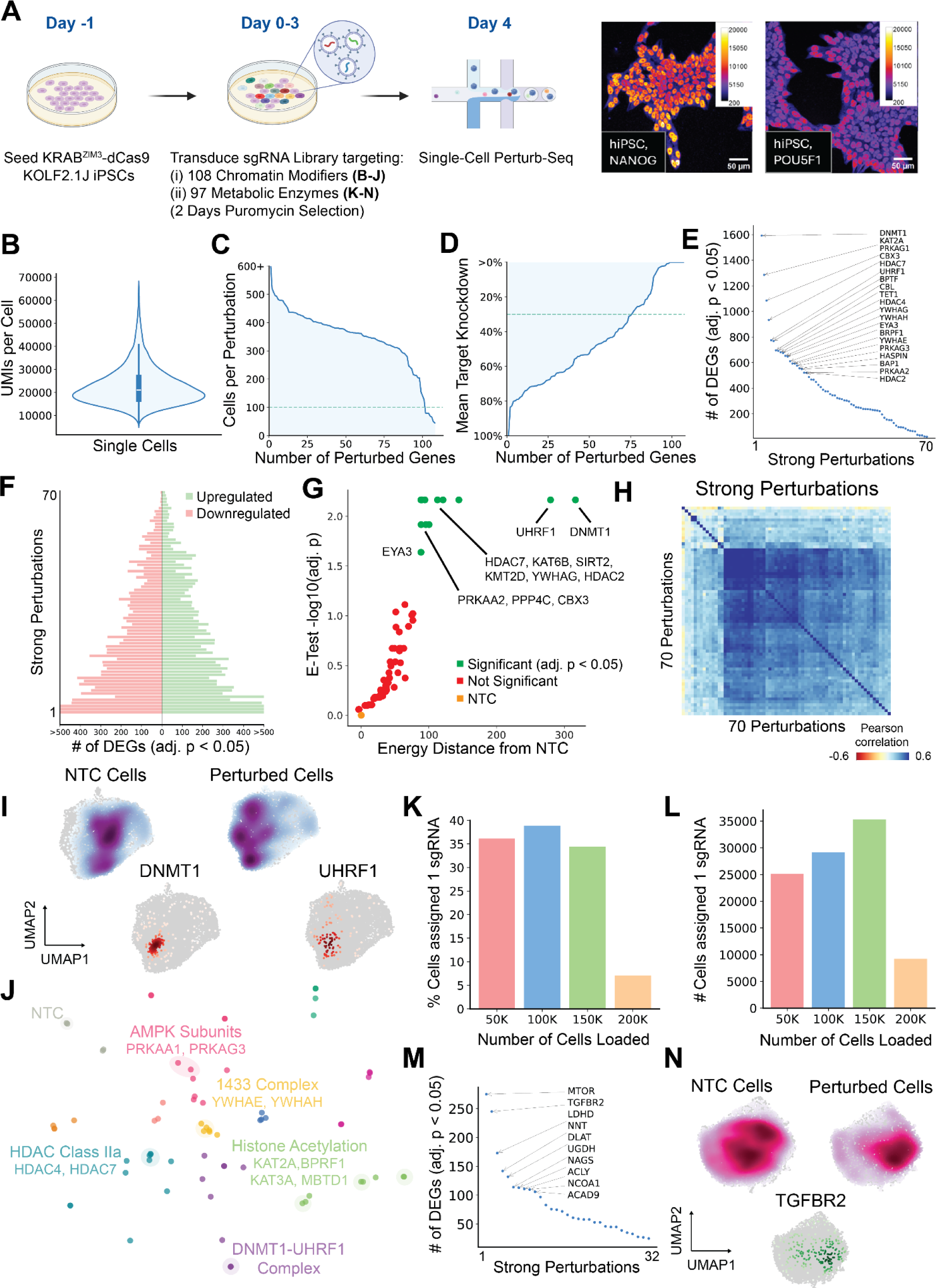
Pilot Perturb-Seq Screens of Epigenome Modifiers and Metabolic Enzymes in Undifferentiated hiPSCs. **(A)** *In Vitro* Perturb-Seq pilot screens experimental scheme. KOLF2.1J iPSCs constitutively expressing KRAB^ZIM3^-dCas9 were transduced with a sgRNA library targeting two libraries: (1) a panel of 108 chromatin modifying genes and (2) a panel of 97 genes involved in metabolism. Following 2 day recovery, cells receiving perturbations were selected via puromycin for 2 days. On day 4, cells were harvested for single cell RNA sequencing. Immunostaining of pluripotency factors NANOG and POU5F1 (OCT4) on day 0 is shown. **(B-J)** are results from screen (1), **(K-N)** are results from screen (2). **(B,C,D)** Chromatin Modifiers Perturb-Seq dataset quality control metrics. **(E)** Number of DEGs per strong perturbation (defined as perturbations passing our filtering metrics (see Methods) and having >= 25 DEGs), with the top 20 perturbations labeled. **(F)** Number of up- and down-regulated DEGs per strong perturbation. **(G)** Energy distance permutation testing (10,000 permutations) of strong perturbations vs NTC. **(H)** Pearson correlation clustermap of pseudo bulked transcriptional profiles of strong perturbations. **(I)** Cell-level UMAP of strong perturbations colored by NTC cell density and perturbed cell density with vignettes of DNMT1 and UHRF1 perturbations. **(J)** Minimum Distortion Embedding (MDE) of pseudo bulked transcriptional profiles of strong perturbations. Clusters are labeled with associated enriched GO/CORUM terms. **(K,L)** Effect of loading variable number of cells per 10X channel on yield of cells with 1 sgRNA assigned for Metabolic Enzymes Perturb-Seq dataset. Aggregation was only performed on channels with 50K, 100K, and 150K loaded cells. **(M)** Number of DEGs per strong perturbation (defined as perturbations passing our filtering metrics (see Methods) and having >= 25 DEGs), with the top 10 perturbations labeled. **(N)** Cell-level UMAP of strong perturbations colored by NTC cell density and perturbed cell density with vignette of TGFBR2 perturbation.

**Supplementary Figure 2:**
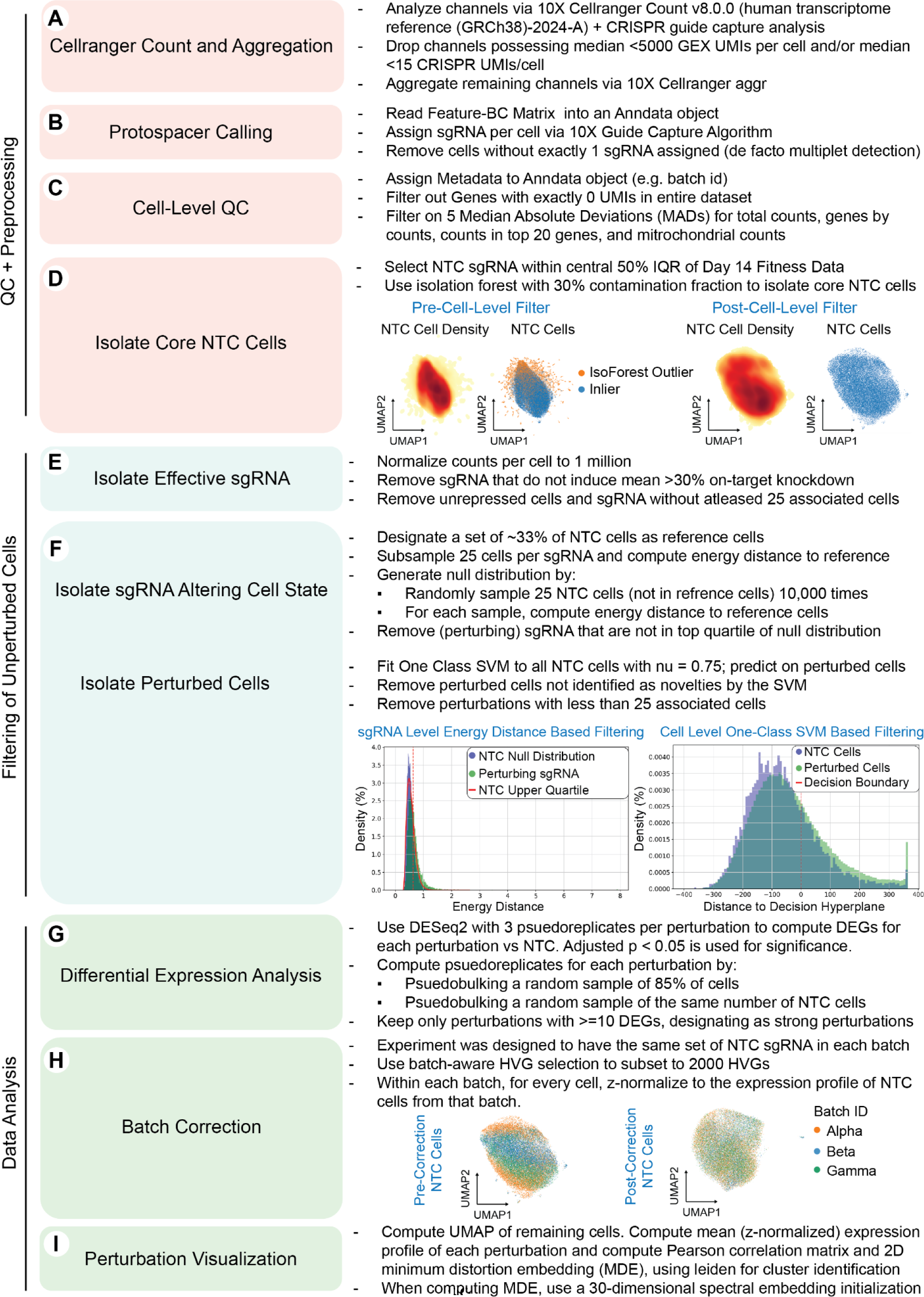
Analysis Pipeline for hiSPC Perturbation Atlas Dataset. **(A-D)** Preprocessing/QC steps to isolate single, live cells with a single sgRNA assigned and a population of core NTC cells. **(E,F)** Filtering steps to identify sgRNA inducing phenotypic change and cells experiencing phenotypic change **(G-I)** Batch correction and data analysis steps to visualize perturbations inducing similar phenotypes.

**Supplementary Figure 3:**
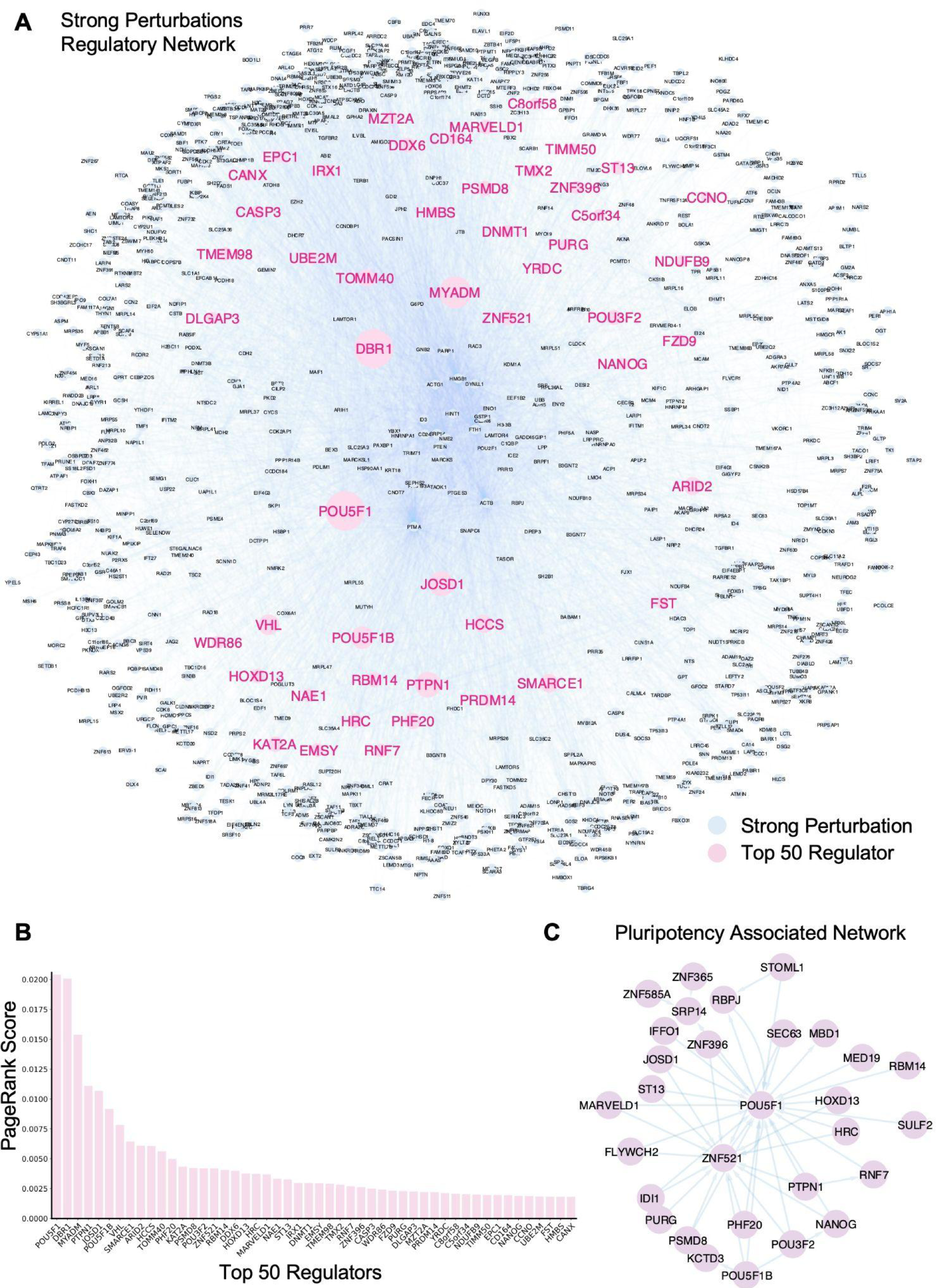
Strong Perturbation Regulatory Network. **(A)** A graphical representation of the regulatory interactions between strong perturbations. Each strong perturbation is a node, and a directed outgoing edge is drawn from a node and every other node that is a DEG of that node (i.e. is regulated by it). Node sizes correspond to their PageRank node importance, and the Top 50 Regulators (as identified by pagerank) are highlighted in pink. **(B)** Top 50 regulators as identified by PageRank. **(C)** Regulatory interactions between perturbations in MDE clusters related to pluripotency.

**Supplementary Figure 4:**
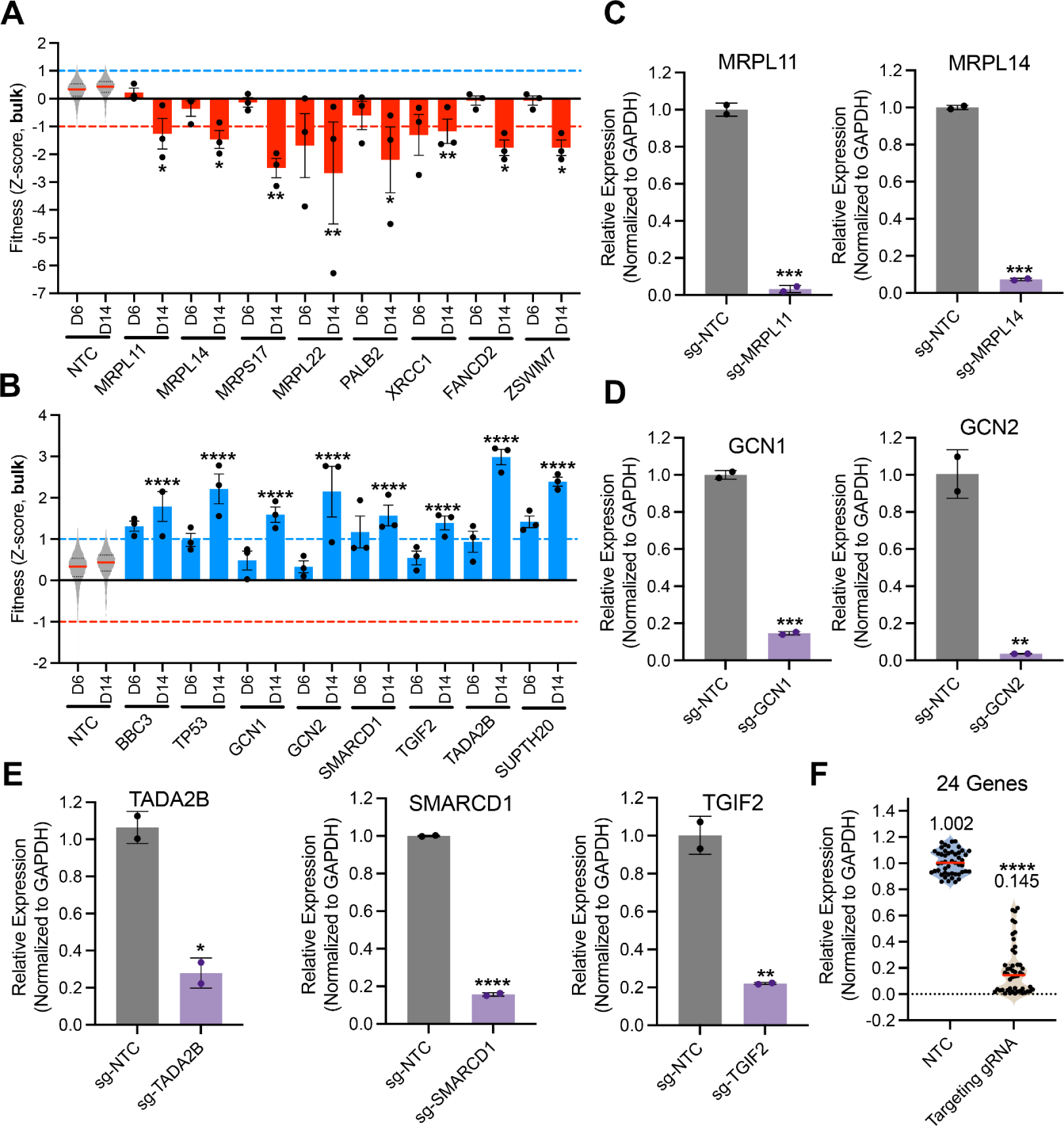
Undifferentiated hiPSC Fitness Screens. **(A, B)** Fitness Z-score at Day6 (D6) and day 14 (D14) in KOLF2.1J KRAB^ZIM3^-dCas9 expressing cells at the perturbation level. Each dot represents a different gRNA, 3 gRNA/perturbation and 478 gRNAs for NTC conditions. Red bars are perturbations with z-score<-1 at day 14, blue bars are perturbations with z-score>1 at day 14. Error Bars represent SD and asterisks indicate significant differences compared to sg-NTC condition (*p < 0.05, **p < 0.01, ***p < 0.001, ****p < 0.0001) **(C, D, E,F)** mRNA gene expression assessed by qPCR between NTC (sg-NTC) and targeting gRNA (sg-Gene) in transduced KRAB^ZIM3^-dCas9 KOLF2.1J. Quantified mRNA are labeled on top of the bar graph. Error bars represent SD and asterisks indicate significant differences (*p < 0.05, **p < 0.01, ***p < 0.001, ****p < 0.0001).

**Supplementary Figure 5:**
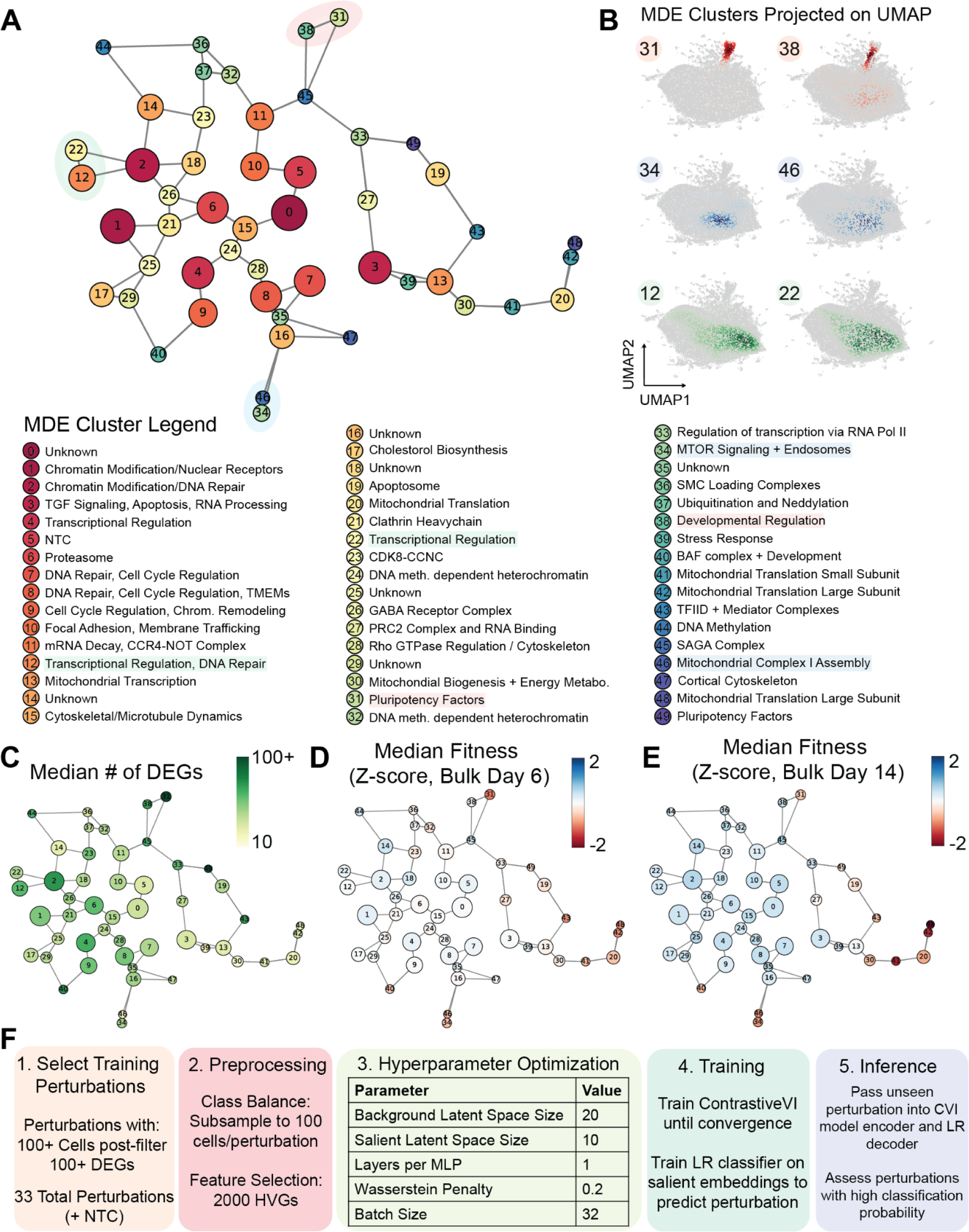
Visualization of Strong Perturbations and Model Training Loop. **(A)** Perturbation similarity MDE map of KOLF 2.1J hiPSCs derived from Figure 2C. Each node represents cluster centroids, with the size of the node corresponding to the number of cluster members. Nodes are connected by edges to their 2 nearest neighbors. Nodes are labeled based on enriched GO/CORUM terms and GSAI descriptions. **(B)** UMAP projection of adjacent MDE clusters, illustrating the spatial organization of clusters 31, 38, 34, 46, 12, and 22 on a single-cell level. **(C)** MDE map colored by the median number of DEGs within each cluster. Color indicates DEGs count, with green representing higher number of DEGs. **(D, E)** MDE map colored by the median day 6 (D) and day 14 (E) fitness Z-score within each cluster. Blue indicates positive fitness effects, and red indicates negative fitness impacts, with nodes sized based on fitness magnitude. **(F)** Training loop and hyperparameter selections for machine learning model. 33 perturbations were chosen initially as they had a high number of associated cells and strong evidence of inducing a cell state change.

**Supplementary Figure 6.**
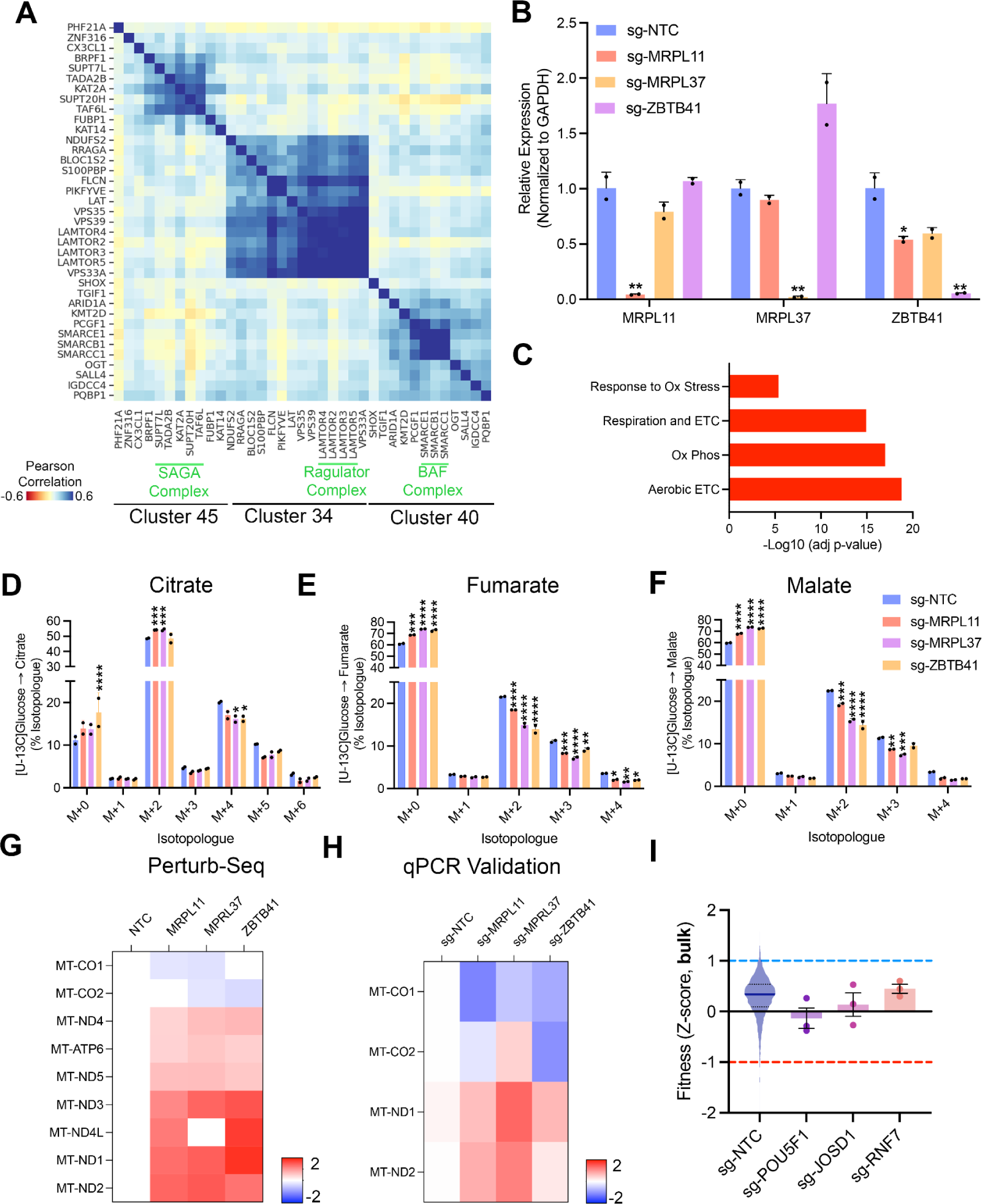
Validation of Known and Novel hiPSC Pluripotency/Mitochondrial Factors. **(A)** Pearson correlation between perturbation MDE clusters with known interactors highlighted in green for cluster 45/SAGA Complex, Cluster 34/Ragulator Complex and Cluster 40/BAF Complex. **(B)** Relative expression levels of MRPL11, MRPL37, and ZBTB41 normalized to GAPDH across different sgRNA treatments. Cells were treated with sgRNAs targeting MRPL11 (orange), MRPL37 (purple), ZBTB41 (yellow), and a non-targeting control (NTC, blue). Error bars represent SD, and asterisks indicate significant differences to sg-NTC condition (*p < 0.05, **p < 0.01). **(C)** Enrichment analysis highlighting significant biological processes associated with MRPL11, MRPL37, and ZBTB41 perturbations. The bar graph displays the negative log-transformed adjusted p-values (-Log10 adj p-value) for pathways including Response to Oxidative Stress, Respiration and Electron Transport Chain (ETC), Oxidative Phosphorylation (Ox Phos), and Aerobic ETC. **(D)** Isotopologue distribution of [U-13C]Glucose into citrate. Bar graph representing the isotopologue distribution (M+0 to M+6) of citrate in cells treated with various sgRNAs (MRPL11, MRPL37, ZBTB41, and NTC). (**E)** Isotopologue distribution of [U-13C]Glucose into fumarate. Bar graph showing the isotopologue distribution (M+0 to M+4) of fumarate across the same sgRNA treatments as panel C. **(F)** Isotopologue distribution of [U-13C]Glucose into malate. Bar graph illustrating the isotopologue distribution (M+0 to M+4) of malate under different sgRNA treatments, with statistical significance denoted (*p < 0.05, **p < 0.01, ***p < 0.001, ****p < 0.0001). **(G)** Perturb-Seq analysis of mitochondrial gene expression (Log2FC) across different sgRNA treatments. Heatmap showing the Log2 fold change of mitochondrial genes in cells transduced with sgRNAs targeting MRPL11, MRPL37, and ZBTB41 compared to sg-NTC. **(H)** qPCR validation of mitochondrial gene expression (Log2FC) across different sgRNA treatments. Heatmap displaying the qPCR validation of mitochondrial gene expression, corresponding to the genes measured in panel B. **(I)** Fitness scores (Z-score) of bulk cells under different sgRNA conditions. Violin plot showing the fitness scores of cells treated with sgRNAs targeting POU5F1, JOSD1, RNF7, and NTC. Error bars indicate SD, with dashed lines representing Z-score thresholds for positive and negative fitness effects.

## METHODS

### Cell Culture

KOLF2.1Js were maintained under feeder-free conditions in mTeSR1 medium (Stem Cell Technologies). Prior to passaging, tissue-culture plates were coated with growth factor-reduced Matrigel (Corning) diluted in DMEM/F-12 medium (Thermo Fisher Scientific) and incubated for 30 minutes at 37 ⁰C, 5% CO2. Cells were dissociated and passaged using the dissociation reagent Versene or Accutase (Thermo Fisher Scientific). HEK293T cells were cultured in Dulbecco’s Modified Eagle Medium (Thermo Fisher Scientific) supplemented with 10% fetal bovine serum (Thermo Fisher Scientific A5256701) and maintained at 37°C in a humidified incubator with 5% CO₂. Cells were passed every 2-3 days upon reaching ∼80% confluence using 0.05% trypsin-EDTA (Thermo Fisher Scientific) for dissociation.

### KOLF2.1J Genome-Scale Perturb-Seq Experiment

We engineered a KOLF2.1J line that stably expresses KRAB^ZIM3^-dCas9^15,22^, and designed a CRISPRi sgRNA library which spans all protein coding genes expressed at > 1.5 RPKM (assayed through bulk RNA-seq) in KOLF2.1Js. In total, 11,739 unique genes were targeted. To process the screen with reasonable workload, we had to split the screen into three different batches. Batch ALPHA contains a library of all annotated DNA binding and modifying enzymes which were determined by combining existing studies and web-scraping of Uniprot. Batch BETA and GAMMA contain all other genes expressed at over 1.5 RPKM, split evenly between the two batches. The total library consists of three sgRNA per gene target (35,217 sgRNA total), and 478 non-targeting control (NTC) sgRNAs, cloned into the CROP-Seq vector ^10^, and delivered to KOLF2.1Js via lentiviral infection at low MOI (<∼0.3) at day 0. Starting at day 2, cells receiving perturbations were enriched via puromycin selection until day 6, at which point cells were harvested for scRNA-Seq and day 6 fitness screening. Remaining cells were allowed to continue propagating to day 14, where they were harvested for day 14 fitness screening. For scRNA-Seq, 115,000 transduced day 6 cells/channels were loaded onto a Next GEM Chip M Single Cell (10x Genomics), for each batch ALPHA, BETA and GAMMA 32 channels across 2 Next GEM Chip M were loaded for a total of 96 channels, samples were processed according to the manufacturer (10x Genomics, User Guide, CG000416). Gene Expression (GEX) libraries and downstream NGS were prepared and run according to the manufacturer Chromium Next GEM Single Cell 3’ HT Reagent Kits v3.1 (10x Genomics).

### RNA Extraction and Quantitative Real-Time PCR

Total RNA was extracted from cells using the RNeasy Mini Kit (QIAGEN) according to the manufacturer’s protocol. For cDNA synthesis, 1000 ng of total RNA was reverse transcribed using the ProtoScript® First Strand cDNA Synthesis Kit (New England BioLabs), following the manufacturer’s instructions. Quantitative real-time PCR (qPCR) was performed using SYBR Green Supermix (Bio-Rad) on a Bio-Rad CFX96 Real-Time PCR Detection System. Gene expression levels were quantified relative to endogenous controls, with each sample measured in technical duplicates. The expression of target genes was normalized to the housekeeping gene GAPDH, and data was analyzed using Bio-Rad CFX Manager software.

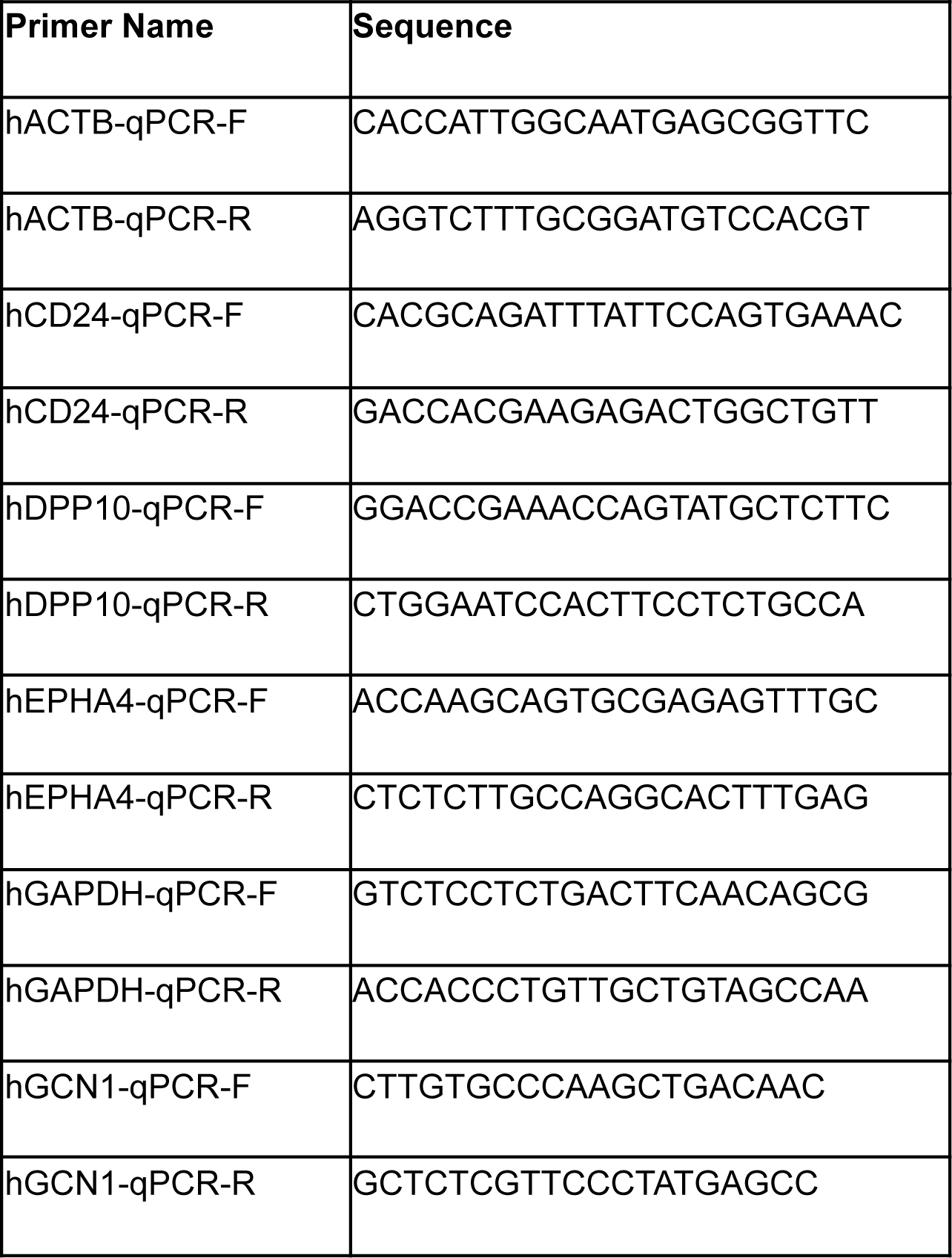
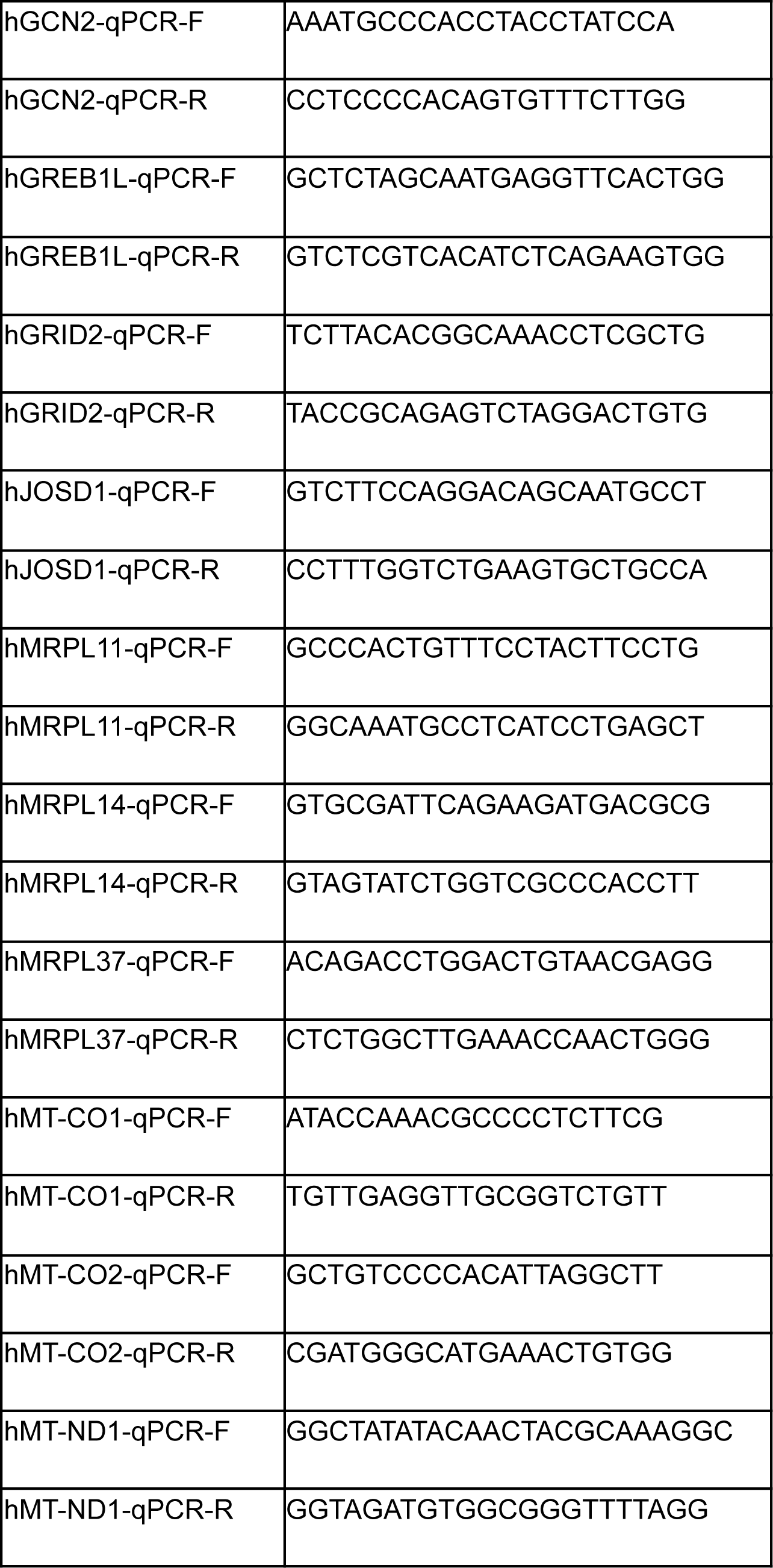
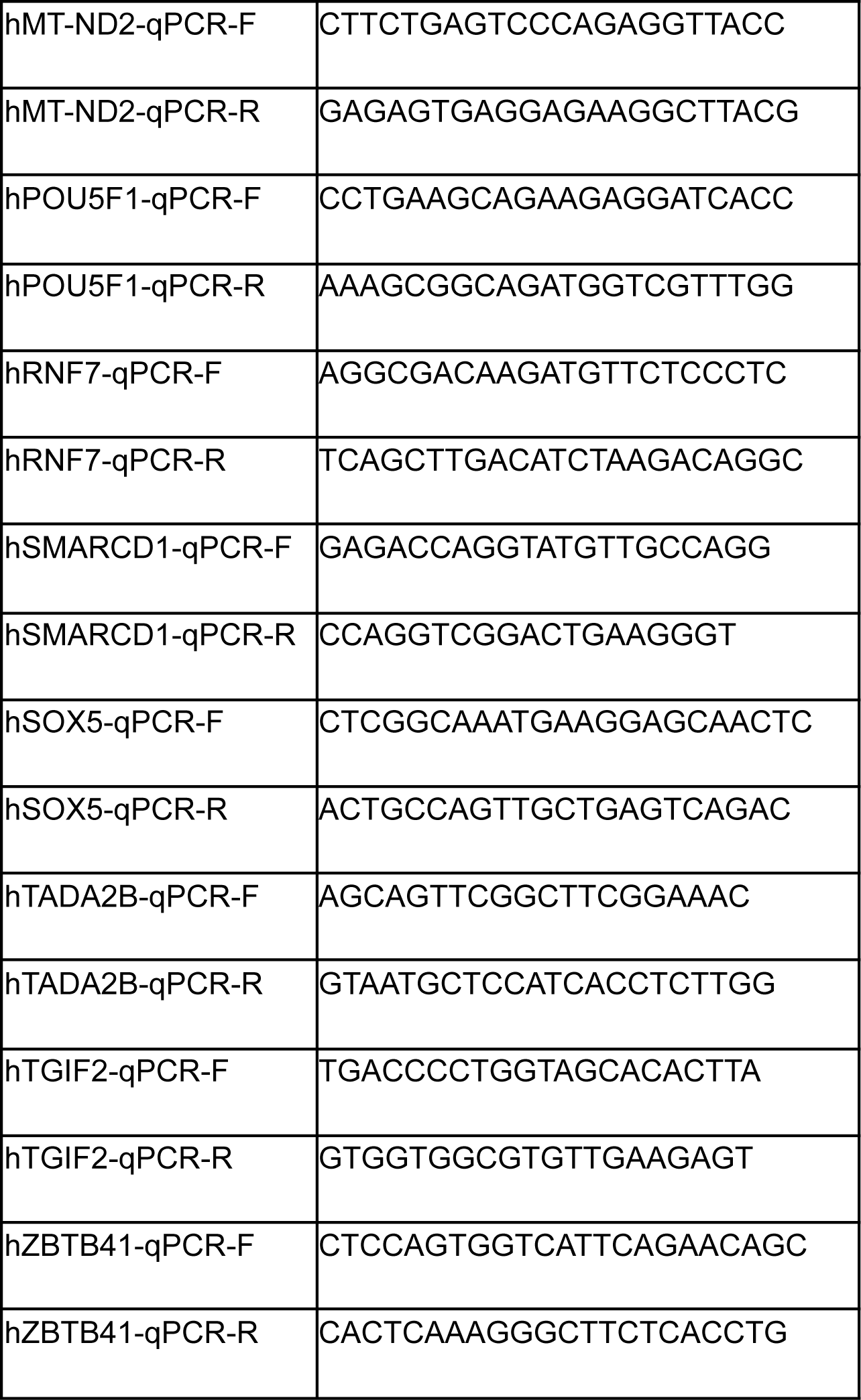

### Immunostaining and microscopy

After a wash with 40 µL phosphate buffer saline (PBS), cells were fixed with 40 µL 4% paraformaldehyde (Thermo Scientific Chemicals 043368.9M) in PBS for 15 minutes. Cells were permeabilized with 40 µL 0.1% Triton-X100 (Sigma T8787) in 3 consecutive washes, 5 minutes each, and washed once with 40 µL PBS. Afterwards, cells were incubated with rabbit Human Protein Atlas antibodies (HPA072117 for NANOG, HPA058267 for POU5F1) at 2 µg/mL and mouse anti-tubulin antibodies (Abcam ab7291) at 1 µg/mL, diluted in PBS + 4% FBS, at room temperature for 2 hours. The samples were washed with 40 µL PBS, 4 × 10 minutes. After the wash, the samples were incubated with secondary antibodies: goat-anti-rabbit IgG, conjugated with Alexa Fluor 488 (Invitrogen A11034), and goat-anti-mouse IgG1, conjugated with Alexa Fluor 555 (Invitrogen A21127), at 1.25 µg/mL in PBS + 4% FBS for 90 minutes at room temperature. Cells were counterstained with DAPI (Invitrogen 62248) in PBS at 1.25 µg/mL for 10 minutes, then washed with 40 µL PBS, 4 × 10 minutes and kept in PBS for imaging. The cells were imaged with Operetta CLS high-content analysis platform (Revvity) at 20x magnification (water immersion, NA 1.0). For each field of view, a z-stack of 17 planes set at a distance of 0.8 µm was collected. The z-stacks were processed in FIJI to obtain a maximum projection image.

### SEC-MS

SEC-MS was performed as previously described by Fossati et al^89^. Briefly, KOLF2 cells were lysed in HNN buffer (50mM HEPES pH7, 150mM NaCl) supplemented with 0.5% NP-40 and protease and phosphatase inhibitor. After sonication to shred DNA, samples were cleared by ultracentifugation (40.000g for 15min at 4C) and 1mg of protein lysate (1mg/ml) was loaded on a 30 MWCO Amicon column (Millipore, UFC803096) to dilute NP-40 with 3 washes of HNN buffer followed by concentration to a volume of 110μL that was injected onto an HPLC system (Agilent, 1260 Infinity II LC system) connected to a BioSep SEC-s4000 size exclusion column (Phenomenex). Each SEC run was performed using a flow of 0.5ml/min of HNN buffer and 5 fractions/min were collected 8.5min after sample loading for a total run time of 26min. Fraction #7 to 78 (72 fraction total) were further processed to be acquired by MS. Proteins in fractions were reduced by adding 100μL of reduction buffer (8M Urea (4M final), 50mM HEPES pH7 and 10mM DTT (5mM final)) and incubated at 37C for 45min before aklylation by addition of 10mM final of iodoaceta and incubated 30 minutes in the dark. Fractions were further diluted with 800μL of 50mM HEPES and incubated overnight after addition of 1μg of Trypsin (Promega, V511X). The next day samples were desaled with C18 96 well plates according to manufacturer instructions and illusions were vacuum dried and stored at −80 °C until MS acquisition. Fractions were resuspended in 60μL of 0.1% formic acid and 5μL of 3x 24 consecutive fractions were pooled to generate an ion mobility gas-phase fractionation (GPF) library was acquired based on the work of Penny et al^90^. Briefly, data was collected from 400 to 1200 m/z using 5 m/z windows with a 1 m/z overlap on each side. The ion mobility range spanned 0.57–1.47 V/cm2, with two quadrupole positions per ion mobility cycle. These windows were evenly distributed across seven acquisition methods, with each method comprising 15 ion mobility cycles. For the SEC fractions, The peptides were sprayed through a 20 mm ZDV emitter kept at 1600 V and 200 degrees C. The mass spectrometer was operated in positive mode with a TIMS range 0.60 Vs/cm2 to 1.60 Vs/cm2. MS1 scans were collected from 100-1700 m/z, while MS2 scan were collected using DIA-PASEF acquisition^91^ (PMID: 33257825) with a 303-1200 m/z range comprised of 24 variable MS2 windows^92^ (PMID: 35944843). Analysis of spectra were performed using Spectronaut^93^ (version 19.2.240905.62635) using default settings. Spectronaut protein-group summarized intensities were used for all downstream SEC-MS analyses.

The full SEC-MS data matrix (8,251 proteins X 72 fractions) for each of three samples were column-normalized using a robust local-regression method to fit a smooth curve through the global intensity trends across each sample. The goal was to detect and preserve global protein intensity trends across fractions as a smooth curve, and adjust protein intensities to this global trend to account for run-to-run noise in MS recovery efficiency. The first step in normalization was to calculate a summary intensity value per fraction as the column effect from a median polish procedure, using the R function medpolish, on the full log-transformed intensity matrix. Next, cubic polynomial curves were fitted to the summary data using exhaustive sliding windows 20 fractions wide with function rlm from the MASS package of R. The residuals of each fraction to all of the local-regression curves that include the fraction were pooled and the median taken as each fraction’s normalization factor. All protein intensities in each fraction were then adjusted by the corresponding fraction’s normalization factor by subtraction on log transformed data. Protein intensity values were then transformed back to their non-log-transformed values, and scaled per protein and per sample by dividing by the sum of each protein’s intensity in a sample. To calculate a combined cosine similarity between protein elution profiles across the three replicates of 72 fractions each, the normalized matrices were concatenated fraction-wise into a global matrix of 8,251 proteins by 216 fractions. Missing values were set to 0. Cosine similarities were then calculated between all possible protein pairs as the similarity between vectors of length 216.

Enrichment of physical interactions detected by SEC-MS in Perturb-Seq derived clusters was calculated by a comparison of the 95th percentile score of within-cluster cosine similarities compared with a random distribution of 95th percentiles calculated on 10,000 permuted clusterings in the same data. Each permutation shuffled the gene labels in the cluster table so that cluster sizes remained the same per cluster name, and the score of each cluster was compared against permutations on the same-named cluster. The portion of permutations with 95th percentile score greater than or equal to the original cluster was used as the p value, p values were adjusted for multiple testing across the 51 clusters by the Benjamini-Hochberg FDR method, and those clusters with FDR < 0.05 were considered significantly enriched.

### Metabolic tracing/Mitochondrial Flux

For isotopic labeling experiments, cells were cultured in mTESR containing 17.5 mM [U13C6]glucose (Cambridge Isotope Laboratories, Inc.) for 24 hours prior to extraction. To extract metabolites, cells were rinsed with saline solution. Then 700 uL of ice-cold 60:40 methanol:water solution was added to the cells along with polar and fatty acid internal standards. Cells were scraped from the plate, and lysates were transferred to Eppendorf tubes. Samples were vortexed for 5 minutes, and 10% was taken for protein quantification by BCA. The remaining homogenate was combined with 500uL of chloroform, vortexed for 5 minutes, and centrifuged (5 min, 21000g, 4C). The upper polar phase was collected, dried under a vacuum at 4C, and derivatized with 2% (w/v) methoxyamine hydrochloride (Thermo Fisher Scientific) in pyridine for 60 min followed by sialyation in N-tert-butyldimethylsilyl-N-methyltrifluoroacetamide (MTBSTFA) with 1% tert-butyldimethylchlorosilane (tBDMS) (Regis Technologies, Inc.) at 37 °C for 30 min. The polar phase was analyzed by gas chromatography (GC)-mass spectrometry (MS) as previously described ^94^ using a DB-35MS column (30 m × 0.25 mm i.d. x 0.25 μm, Agilent J&W Scientific) installed in an Agilent 7890A GC interfaced with an Agilent 5975C MS.

### Guide Library Generation

To target genes by CRISPRi, we selected the top3 gRNAs/genes from the CRISPick tool^23,95^. In total we generated 35217 gRNAs targeting genes and 478 Non-targeting gRNAs. This pool of gRNAs has been synthesized in three oligonucleotides pools ALPHA, BETA, and GAMMA (Genescript) and has been cloned into the CROP-Seq vector as follows.

The Gibson assembly reactions were set up as follows: 1000 ng digested backbone, 1:20 molar ratio of insert, 2X Gibson assembly master mix (New England Biolabs), H20 up to 100 μl. After incubation at 50 °C for 1 h, the product was transformed via electroporation into One Shot Stbl4 Electromax chemically competent Escherichia coli (Invitrogen). A fraction (1 µL) of cultures was spread on carbenicillin (50 µg/ml) LB plates and incubated overnight at 37 ⁰C to estimate coverage, while the rest of the transformant were seeded into 150 ml of LB medium containing carbenicillin (50 µg/ml) and incubating overnight in a shaker at 37 ⁰C for 16-18 h. The plasmid DNA was then extracted using a Plasmid Maxi Kit (Qiagen).

sgRNAs used for validation experiments

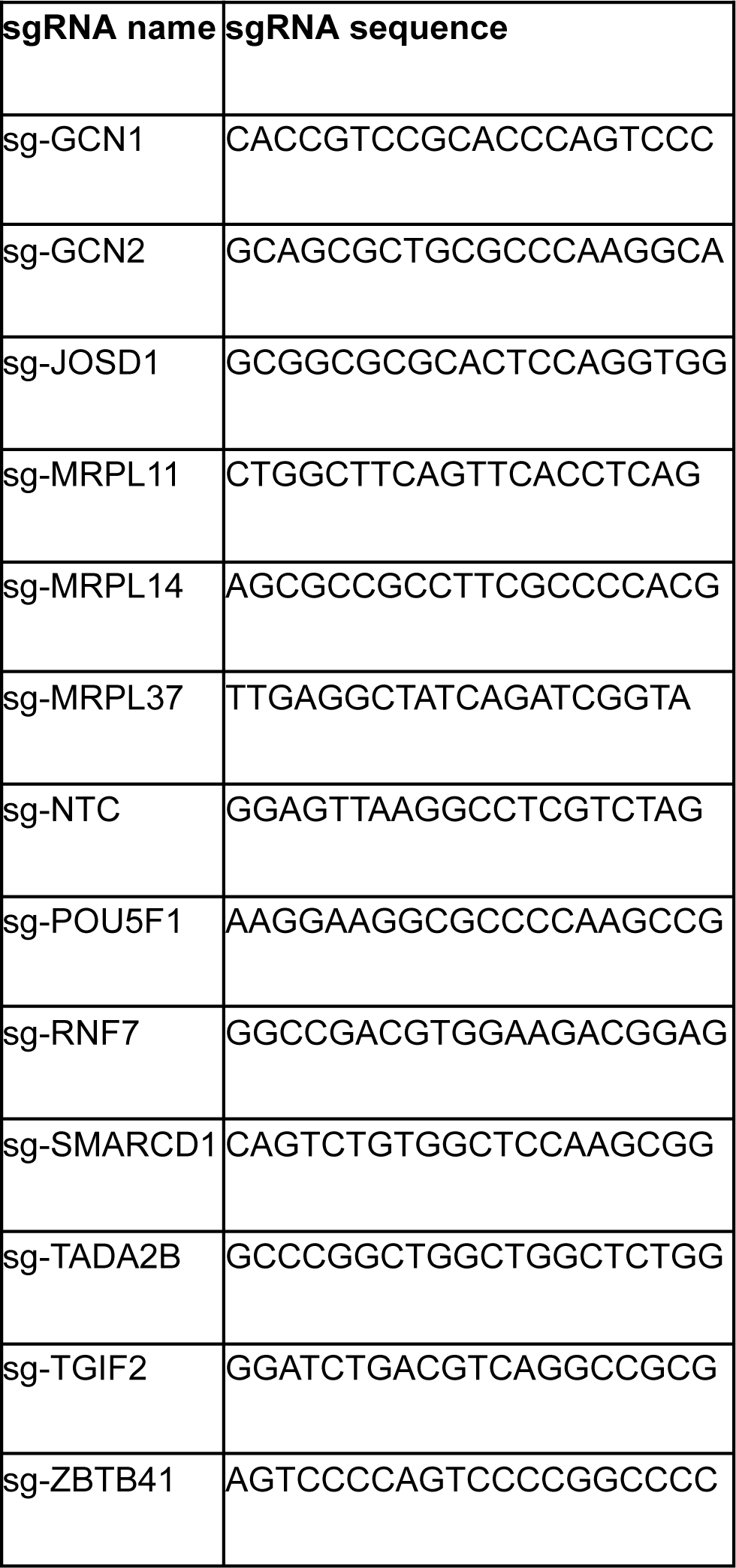

### Library Lentiviral Production

HEK 293T cells were maintained in high glucose DMEM supplemented with 10% fetal bovine serum. Cells were seeded in a 15 cm plate 1 day prior to transfection, such that they were 60–70% confluent at the time of transfection. For each plate, 36 μl of Lipofectamine 2000 (Thermo Fisher Scientific 11668027) was added to 2 mL of Opti-MEM (Thermo Fisher Scientific 31985062). Separately 3 μg of pMD2.G (Addgene #12259), 12 μg of pCMV delta R8.2 (Addgene #12263) and 9 μg of the pooled gRNA vector library was added to 2 mL of Opti-MEM. After 5 minutes of incubation at room temperature, the Lipofectamine 2000 and DNA solutions were mixed and incubated at room temperature for 30 minutes. During the incubation period, medium in each plate was replaced with 20 ml of fresh, pre-warmed medium per well. After the incubation period, the mixture was added dropwise to each plate of HEK 293T cells. Supernatant containing the viral particles was harvested after 48 and 72 hours, filtered with 0.45 μm filters (Steriflip, Millipore), and further concentrated using Amicon Ultra-15 centrifugal ultrafilters with a 100,000 NMWL cutoff (Millipore) down to a final volume of 600–800 μl, for each 15 cm plate. Finally, the concentrated supernatant was divided into 200 μl aliquots and frozen at −80°C. Two 15 cm plates worth of lentivirus was adequate.

### gRNA amplification

gRNA barcodes were amplified from cDNA generated by the 10x single cell platform, and gDNA from cells harvested at day 6 and 14 of the KOLK2.1J fitness screen. Amplicons were prepared for deep sequencing through a two-step PCR process.

For amplification of barcodes from cDNA, the first step was performed as four separate 50 μl reactions for each sample. 2.5 μl of the cDNA was input per reaction with Kapa Hifi Hotstart ReadyMix (Kapa Biosystems KK2602). The PCR primers used were, CROPseq_sgRNA_barcodeAmp_Fprimer: GACTGGAGTTCAGACGTGTGCTCTTCCGATCTCTTGTGGAAAGGACGAAACAC and NEBNext Universal_R: GACTGGAGTTCAGACGTGTGCTCTTCCGATCTCTTGTGGAAAGGACGAAACAC. The thermocycling parameters were 95°C for 3 min; 20–26 cycles of (98°C for 20 s; 65°C for 15 s; and 72°C for 30 s); and a final extension of 72°C for 5 min. The numbers of cycles were tested to ensure that they fell within the linear phase of amplification. Amplicons (∼450 bp) of 4 reactions for each sample were pooled, size-selected and purified with Agencourt AMPure XP beads (Beckman Coulter, Inc.) at a 0.8 ratio. The second step of PCR was performed with two separate 50 μl reactions with 50 ng of first step purified PCR product per reaction. NEBNext Multiplex Oligos for Illumina (Dual Index Primers) were used to attach Illumina adapters and indices to the samples. The thermocycling parameters were: 95°C for 3 min; 6 cycles of (98°C for 20 s; 65°C for 15 s; 72°C for 30 s); and 72°C for 5 min. The amplicons from these two reactions for each sample were pooled, size-selected and purified with Agencourt AMPure XP beads at an 0.8 ratio. The purified second-step PCR library was quantified by Qubit dsDNA HS assay (Thermo Fisher Scientific) and used for downstream sequencing on an Illumina HiSeq platform.

For amplification of barcodes from genomic DNA, genomic DNA was extracted from stored cell pellets with a DNeasy Blood and Tissue Kit (Qiagen). The first step PCR was performed as two separate 50 μl reactions for each sample. 500 ng of genomic DNA was input per reaction with Kapa Hifi Hotstart ReadyMix. The PCR primers used were, sgRNAsequenceAmp_gDNA_F: ACACTCTTTCCCTACACGACGCTCTTCCGATCTTATATATCTTGTGGAAGGACGAAACACCG and sgRNAsequenceAmp_gDNA_R: GACTGGAGTTCAGACGTGTGCTCTTCCGATCTCCTTATTTTAACTTGCTATTTCTAGCTCTA. The thermocycling parameters were: 95°C for 3 min; 24–32 cycles of (98°C for 20 s; 60°C for 15 s; and 72°C for 30 s); and a final extension of 72°C for 5 min. The numbers of cycles were tested to ensure that they fell within the linear phase of amplification. Amplicons (∼150 bp) of the two reactions for each sample were pooled, size-selected with Agencourt AMPure XP beads at a ratio of 1.6. The second step of PCR was performed as two separate 50 μl reactions with 25 ng of first step purified PCR product per reaction. NEBNext Multiplex Oligos for Illumina (Dual Index Primers) were used to attach Illumina adapters and indices to the samples. The thermocycling parameters were: 95°C for 3 min; 6–8 cycles of (98°C for 20 s; 65°C for 20 s; 72°C for 30 s); and 72°C for 2 min. The amplicons from these two reactions for each sample were pooled, size-selected with Agencourt AMPure XP beads at a ratio of 1.6. The purified second-step PCR library was quantified by Qubit dsDNA HS assay (Thermo Fisher Scientific) and used for downstream sequencing on an Illumina NovaSeq platform.

### Fitness Screen Analysis

CROP-Seq plasmid, Day6, and Day14 gRNA amplicon libraries (ALPHA, BETA, GAMMA) were sequenced on NovaSeq X Sequencer (Illumina). FATSQs files were analyzed using the MAGeCK python package for computing enrichment and depletion to the plasmid libraries at the gRNA and gene levels^96^. Fitness score correspond to a Z-score was calculated as followed:

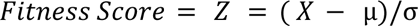

Where X is Log2Fold change, µ is the mean of all Log2FoldChange, σ is the standard deviation of all Log2fold change.

### Statistical Analysis

All statistical analyses for validation experiments were performed using GraphPad Prism 9. For **Figures 3F**, **Figure 6, and Supplementary Figure 6**, statistical comparisons between conditions were conducted using a two-way ANOVA with multiple comparisons against the non-targeting control (NTC). For qPCR experiments in **Supplementary Figure S4 and Figure 6**, comparisons were made using a two-tailed unpaired t-test against the NTC control condition. Statistical details, including exact p-values and significance levels, are provided in the results, figures, and figure legends.

### Perturb-Seq Analysis Pipeline

We developed a data analysis pipeline for analyzing the single-cell Perturb-Seq data set established in this study The analysis pipeline can be openly accessed at: github link and is described below as well as **Supplementary Figure 2**.

### Genome Reference Alignment, Cell Calling, sgRNA Assignment, and Aggregation

Genome reference alignment, cell calling, sgRNA assignment, and channel aggregation were all performed using cellranger v8.0.0 count and aggr using the cellranger human transcriptome reference (GRCh38)-2024-A and default command line arguments. Aggregation was performed on all channels passing basic QC (see Supplementary) with >5000 median GEX UMIs per cell and >15 median CRISPR UMIs per cell (corresponding to 90/96 channels) to ensure sufficient read depth and sgRNA call confidence for all channels with the –no-secondary flag. Aggregation was manually terminated after SC_RNA_AGGREGATOR.WRITE_MATRICES and SC_RNA_AGGREGATOR._CRISPR_ANALYZER.CALL_PROTOSPACERS chunks were completed.

### Metadata Assignment and Cell-Level Quality Control Metrics

The analysis pipeline works directly on the output of cellranger aggr. First, we read the filtered_feature_bc_matrix.mtx.gz data matrix generated by cellranger via anndata .read_h5ad() into an Anndata object. For each cell, the corresponding sgRNA is assigned via the protospacer_calls_per_cell.csv file produced as cellranger aggr output. Cells without exactly 1 sgRNA assigned are removed. We hypothesize that cells with 0 sgRNA assigned (∼25% of all cells) truly do contain a sgRNA as they survived 4 days of puromycin selection, and thus these are cells where the sgRNA simply failed to detect. This may be due to challenges with robustly capturing polyadenylated, linear sgRNA via dial-out PCR. Further, we hypothesize that cells with >1 sgRNA assigned are likely multiplets (two or more cells within a single GEM) as the experimental MOI of < 0.3 would make events where there is >1 sgRNA per cell unlikely. We anecdotally remark that the percent of cells identified with more than 1 sgRNA (∼28%) detected correspond similarly to 10x’s reported approximate multiplet rate based on a targeted recovery of 60,000 single cells. Given our experimental design, we note a data loss of ∼50% of cells at this stage, and advise researchers to bear this loss in mind as a key consideration when determining the cell coverage required for future similar experiments.

After isolating cells with exactly 1 sgRNA assigned, we assign metadata per cell (Run ID, 10X Chip ID, 10X Channel ID, Cell Type (KOLF2.1J), Perturbation Type (CRISPRi)). Genes are removed with exactly 0 UMIs detectable in the entire dataset. Dead and aberrant cells are removed via the median absolute deviation (MAD) method from Heumos et al. (^97^), filtering on 5 MADs for the QC covariates: log1p_total_counts, lop1p_n_genes_by_counts, pct_counts_in_top_20_genes, pct_counts_mt evaluated by scanpy’s sc.pp.calculate_qc_metrics() with cumulative proportion to the 20th most expressed gene (percent_top=[20]).

### Core Control Cell Identification

Although NTC sgRNA are designed to not directly target any genomic loci, there are instances where a NTC sgRNA, perchance, may induce a phenotypic alteration as outlined by ^15^. In light of this, we sought to remove NTC sgRNA and NTC cells that demonstrated phenotypic deviation from the norm, with the underlying assumption that NTC sgRNA typically do not induce phenotypic alterations compared to wild-type KOLF2.1J hiPSCs. To do so, we leveraged the results of fitness screening, removing any NTC sgRNA that was not within the middle 50% interquartile range of fitness log2 fold changes (L2FCs). This corresponded to a NTC sgRNA “whitelist” of 236 sgRNA with mean fitness L2FC of 0.3, max L2FC of 1.0, and min L2FC of −0.4. NTC sgRNAs in the aggregated anndata object were subset down to contain only those in the whitelist. We then used an outlier detection model (isolation forest ^98^) to filter out NTC cells with a deviant phenotype from the norm. NTC cells were isolated from the perturbed cells, batch-aware highly variable gene (HVG) selection was performed using sc.pp.highly_variable_genes with flavor=’seurat_v3’ to select 2000 HVGs, embedded via the top-100 principal components, and filtered using an isolation forest (sklearn) with a 30% contamination fraction. Remaining cells were identified as core NTC cells, and merged back with the perturbed cells.

### sgRNA and Cell Level Quality Control and Filtering

After core NTC detection, counts per cell were normalized to 1 million. The following were filtered out: (1) sgRNA that did not induce, on average over all associated cells, >30% on-target knockdown, (2) sgRNA without at least 25 associated cells, and (3) single cells that did not experience any on-target knockdown. Target knockdown was evaluated relative to the mean target expression within NTC cells from the same batch. Further, we aimed to filter out sgRNA that did not induce a phenotypic effect. For each batch, a random subset of 3000 NTC cells were designated as reference cells. The energy distance was then calculated for each perturbing sgRNA to the reference cells (from the same batch) following the procedure outlined in ^18^. Briefly, each perturbing sgRNA was first randomly downsampled to 25 cells. Then, top 2000 highly variable genes were computed and after transformation via log1p and scaling (zero-centering), the top 50 PCs were used as features for each cell. The energy distance was computed between each sgRNA and the reference cells. Additionally, the energy distance was computed between 10,000 random samples of 25 NTC cells (not in the reference cells) and reference cells to create a null distribution. Perturbing sgRNA not within the top quartile of NTC energy distances were filtered out. Next, for each batch, a One-Class Support Vector Machine (SVM) classifier was fitted to all NTC cells with a nu=0.75 (mimicking the same upper-quartile cutoff) to perform novelty detection on perturbed cells from the same batch (using again the top 50 PCs as features). Perturbed cells that were classified as NTC cells were filtered out, indicating cells that are potentially escaping perturbation or not perturbed to a sufficient magnitude. Finally, all perturbations without at least 25 total cells (combining cells from all sgRNA targeting the same gene) are filtered out.

### Differential Expression Analysis

To determine DEGs we leveraged DESeq2 as per the recommendation by Peidli et al. ^18^. As DESeq2 is inherently a bulk RNA seq modality, we generated three pseudoreplicates as suggested by Hafemiester and Halbritter^99^, using 85% of cells associated with each perturbation to generate pseudo replicates, and choosing at random the same number of NTC cells to form reference pseudoreplicates. Perturbed cells are compared to NTC cells from the same batch. Up- and down-regulated perturbations are determined by the sign of the log2 fold change reported by DESeq2. While we did observe at read depths of ∼20,000 UMIs a balance between the number of up- and down-regulated genes (Supplementary Figure 2F), we did notice a bias towards detecting downregulated genes at read depths of ∼5000 UMIs, likely due to dropout events. We feel that determining the optimal methodology for differential expression analysis in Perturb-Seq analysis is still an area of open research. Perturbations with <10 DEGs (expressed genome scale study) or <25 DEGs (pilot screens) were filtered out. Pathways and processes enrichment analysis was performed using Metascape.

### Batch Correction, Visualization, and Clustering

Batch correction was performed using the methodology outlined by Replogle et al.^15^. Counts per cell were normalized to 1 million and log-transformed via sc.pp.log1p. Then, expression profiles for cells within a batch were z-normalized to the mean expression profile of NTC cells from the same batch. That is, 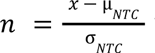 where *n* is the normalized expression profile of a cell, *x* is the unnormalized expression profile of a cell, µ*_NTC_* is the mean expression profile of NTC cells, and σ*_NTC_* is the standard deviation of expression profiles of NTC cells. As in ^15^, we clip normalized expression values to be between [-10,10] to handle extreme outliers. Then, batch-aware HVG selection was performed on raw (non-normalized) counts using sc.pp.highly_variable_genes with flavor=’seurat_v3’ to select 2000 HVGs. Using these 2000 genes as features, two visualizations are generated.

First, the normalized profiles are scaled and embedded via PCA (sc.pp.pca with default parameters). The top 50 PCs are used as features to initialize a neighborhood graph (sc.pp.neighbors with default parameters) and embedded into 2D via UMAP (sc.tl.umap with default parameters). The UMAP embedding density of perturbed cell populations is computed via a kernel density estimate (KDE, sc.tl.embedding_density with default parameters). Clustering is additionally performed using leiden clustering (sc.tl.leiden with default parameters; max_iter=2) Embeddings are visualized using sc.pl.umap and sc.pl.embedding_density.

Second, the pairwise Pearson correlation between pseudo bulked normalized expression profiles of each perturbation is computed (**Figure 2A**). We then embed the correlation matrix into 2-dimensional space using a minimum distortion embedding (MDE). Each pseudo bulk profile is first z-normalized to itself (as in ^15^) to equate the MDE distance optimization problem to a correlation optimization problem. That is, 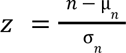 where *z* is the “internally” normalized pseudo bulk expression profile of a perturbation, *n* is the normalized expression profile of a cell, µ*_n_* is the mean value of the normalized expression profile, and σ*_n_* is the standard deviation of the normalized expression profile. Then, we precompute the 30-dimensional spectral embedding of the dataset using n=15 neighbors. This precomputed embedding is used as input to pyMDE’s preserve_neighbors function, where the dataset is embedded into 2D using n=5 neighbors and a repulsive fraction of 15. Clustering is performed using leiden clustering in 2D space using n=8 neighbors. Clusters are annotated via functional enrichment analysis using significant terms output from CORUM complexes and Gene Ontology using python gprofiler package^100^ as well as augmentation using GSAI for unannotated clusters^67^.

### Energy Distance Computation

The energy distance in **Supplementary Figure 1F** is computed using the scperturb python package from ^18^ and associated computational pipeline. Briefly, perturbations are subsampled to 25 cells per perturbation, embedded into 50-dimensional PC space using the top 2000 HVGs as features, and the energy distance is permuted 10,000 times to determine significance.

### Machine Learning Model Architecture and Training

To develop the model in **Figure 4C**, we were inspired by the SATURN architecture^101^, where an unsupervised cell state embedding is passed into a logistic regression model to annotate cell type. We leveraged ContrastiveVI to learn perturbation-specific cell state embeddings. The overall training procedure is outlined in **Supplementary Figure 4F**. We first subset to perturbations with at least 100 cells and 100 associated DEGs, and for class-balancing, we downsample to 100 cells per perturbation using scperturb.equal_subsampling. Initial feature selection was done by using the top 2000 highly variable genes used previously in the MDE/UMAP analysis. Then, we used Raytune/AsyncHyperband for hyperparameter optimization over 500 random trials. Optimal hyperparameter choices are listed in **Supplementary Figure 4F**. We then trained the ContrastiveVI model to convergence with 80/20 test/validation split (negative validation ELBO not improving after 45 epochs) and visualized the salient feature space using UMAP. Finally, we fit a logistic regression classifier (sklearn) using a 80/20 test/train split to the salient features, using as ground truth the known perturbation labels. (**Figure 4E**). For **Figure 4F**, the logistic regression classifier was first fit to the psuedobulked profiles of salient embeddings for each seen perturbation. Then, unseen perturbations were psuedobulked, fed into the ContrastiveVI model encoder, and decoded using the pretrained logistic regression classifier.

